# Distinct upper airway epithelium interferon-stimulated and profibrotic gene expression between adult and infant rhesus macaques infected with SARS-CoV-2

**DOI:** 10.1101/2022.02.12.480218

**Authors:** Stephanie N. Langel, Carolina Garrido, Caroline Phan, Tatianna Travieso, Todd DeMarco, Zhong-Min Ma, Rachel Reader, Katherine J. Olstad, Rebecca L. Sammak, Yashavanth Shaan Lakshmanappa, Jamin W. Roh, Jennifer Watanabe, Jodie Usachenko, Ramya Immareddy, Rachel Pollard, Smita S. Iyer, Sallie Permar, Lisa A. Miller, Koen K.A. Van Rompay, Maria Blasi

## Abstract

The global spread of Severe Acute Respiratory Syndrome Coronavirus 2 (SARS-CoV-2) and its associated coronavirus disease (COVID-19) has led to a pandemic of unprecedented scale. An intriguing feature of the infection is the minimal disease in most children, a demographic at higher risk for respiratory viral diseases. To elucidate age-dependent effects of SARS-CoV-2 pathogenesis, we inoculated two rhesus macaque monkey dam-infant pairs with SARS-CoV-2 and conducted virological and transcriptomic analysis of the respiratory tract and evaluated systemic cytokine and antibody responses. Viral RNA levels in all sampled mucosal secretions were comparable across dam-infant pairs in the respiratory tract. Despite comparable viral loads, adult macaques showed higher IL-6 in serum while CXCL10 was induced in all animals. Both groups mounted neutralizing antibody (nAb) responses, with infants showing a more rapid induction at day 7. Transcriptome analysis of tracheal tissue isolated at day 14 post-infection revealed significant upregulation of multiple interferon-stimulated genes in infants compared to adults. In contrast, a profibrotic transcriptomic signature with genes associated with cilia structure and function, extracellular matrix (ECM) composition and metabolism, coagulation, angiogenesis, and hypoxia was induced in adults compared to infants. Our observations suggest age-dependent differential airway responses to SARS-CoV-2 infection that could explain the distinction in pathogenesis between infants and adults.

## Introduction

The epidemiological evidence has consistently demonstrated that SARS-CoV-2 infections in young children are mostly mild with relatively low hospitalization rates (1, 2). This results in significantly decreased fatality rates of young (0-9 years old) compared with old (>70-year-old) populations (3-9). Despite the increase in pediatric hospitalizations during emergence of the Delta variant, the proportion of children with severe disease after the Delta variant became predominant were similar to those earlier in the pandemic (10). This suggests that the increase in pediatric hospitalizations during this time is related to other factors, and that children are still at significantly lower risk to severe COVID-19 compared to adults.

Differences in local immune responses in the respiratory tract that include antiviral and pro-inflammatory mediators likely play a role in the age-dependent pathogenesis of SARS-CoV-2 (11). A recent study showed that children have increased expression of relevant pattern recognition receptors including MDA5 (*IFIH1*) and RIG-I (*DDX58*) in the upper airways when compared to adults, suggesting that recognition of SARS-CoV-2 entry into respiratory cells is enhanced in children (12). This is relevant for recruitment of fast-acting innate immune cells like neutrophils, which were shown to be higher in children during the acute phase of SARS-CoV-2 infection compared to adults (13). Interferon (IFN) responses are also key determinants of COVID-19 severity, and potent production of type III, and to a lesser extent type I IFNs in the upper respiratory tract are associated with decreased viral load, milder COVID-19 and younger age (14). However, there remains a large gap in our understanding of how age-dependent factors mediate disease severity and recovery post-infection.

Non-human primate (NHP) models are valuable resources to address these questions related to SARS-CoV-2 pathobiology and to explore vaccine and drug-based interventions against COVID-19 (15, 16). In this pilot study, we developed a maternal-infant SARS-CoV-2 infection model to elucidate the age-dependent effects on viral pathogenesis. By using dam-infant pairs and inoculating both dam and infant at the same time, we were able to simultaneously analyze immune responses to SARS-CoV-2 infection in two different age groups while limiting genetic variation. We show that SARS-CoV-2 infection in dam-infant rhesus macaque pairs results in innate and adaptive immune differences including SARS-CoV-2 neutralizing antibody response kinetics and innate immune and profibrotic gene expression in the conducting airways. Our data highlight age-dependent differential immune and lung responses to SARS-CoV-2 infection that may be important to guide the prevention and treatment of severe COVID-19.

## Results

### Experimental design and SARS-CoV-2 infection of dam/infant pairs

To investigate age-dependent differences in pathogenesis of SARS-CoV-2, two dams and their 6-month-old infants were inoculated intranasally and intratracheally with SARS-CoV-2 (USA-WA1/2020 isolate). Nasal and pharyngeal swabs, bronchoalveolar lavage (BAL), blood, breast milk and rectal swabs were collected over time up to day 14 post-challenge (**Fig. 1A**).

**Fig. 1.**
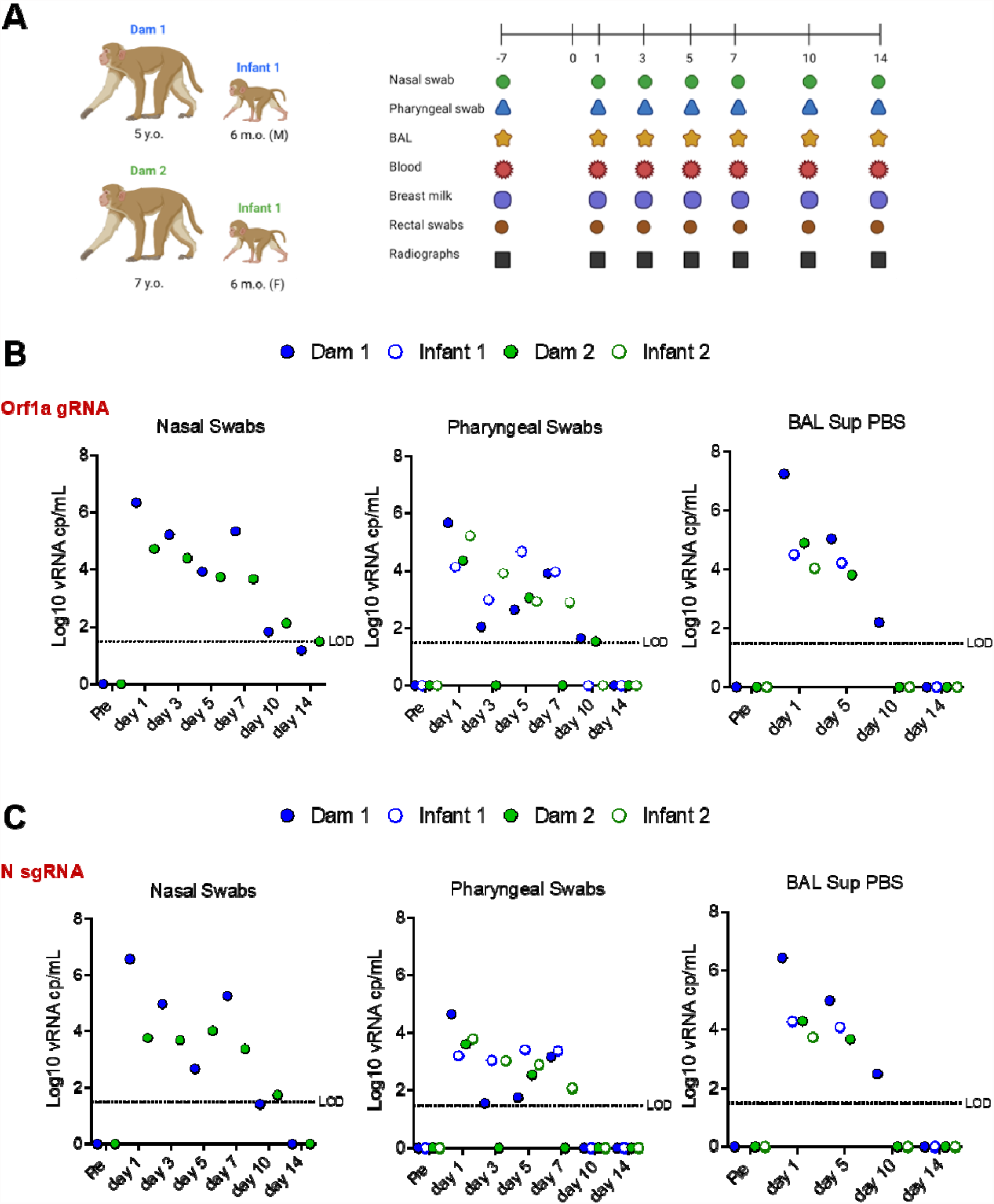
Experimental design and SARS-CoV-2 RNA shedding. **A** Two dams and their 6-month-old infants were inoculated intratracheally and intranasally with 2.5 × 10^6^ PFU and 1.5 × 10^6^ PFU of SARS-CoV-2 (WA strain), respectively. Nasal and pharyngeal swabs, bronchoalveolar lavage (BAL), blood, breast milk and rectal swabs were collected over time up to day 14 post-challenge. Additionally, lung radiographs were performed throughout the experiment. **B** Log_10_ viral RNA copies per mL are reported for nasal swabs, pharyngeal swabs and BAL supernatants over time for both the genomic (Orf1a gene) and **C** subgenomic (N gene) viral RNA.

To compare viral replication kinetics between the adult dams and infant macaques after SARS-CoV-2 infection, we performed both genomic (Orf1a gene) and subgenomic (N gene) PCR on nasal (dams only), pharyngeal, buccal and rectal swabs. Nasal swabs were not collected from infants due to restricted size of infant nostrils. As shown in **Fig. 1B,C**, we did not detect significant differences in viral loads between adults and infants in pharyngeal or BAL supernatant or BAL pellet (**Supplemental Fig. 1**) samples. Low levels of viral RNA were detected in one of the two infants in buccal and rectal swabs, while no viral RNA was detected in breast milk (**Supplemental Fig. 1**).

We evaluated clinical symptoms and radiographic changes overtime. Mild symptoms, including occasional sneezing were reported. The radiographs were scored for the presence of pulmonary infiltrates, according to a standard scoring system (0 to 3 per lobe). Individual lobes were scored and scores per animal per day were totaled. Only the adult macaques had scores on multiple (≥2) days and the highest scores observed (score of 5) were from one of the adult macaques (**Supplementary Table 1**). Overall in both age groups the lesions in the lung were minimal and largely resolved consisting of occasional foci of mild inflammation on HE stained sections (**Supplemental Fig. 2**). Trichrome stained sections did not show an overt increase in fibrosis in either group. There was no significant evidence of interstitial pneumonia in any animal, only occasional focal increase in interstitial cellularity and in some areas an increase in alveolar macrophages, as previously reported (17).

### Systemic innate cellular and cytokine responses in SARS-CoV-2-infected dam and infant rhesus macaques

We evaluated innate immune cell subsets in the peripheral blood and observed no major differences in proportion of cells between infant and dams after SARS-CoV-2 infection (**Fig. 2A**). Interestingly, however circulating neutrophils (lineage-, CD66+) and total HLA-DR+ (lineage-, CD66-) cells, comprising of monocytes and dendritic cell subsets, were higher in adults and infants prior to inoculation with SARS-CoV-2, respectively (**Fig. 2A**). For plasma cytokine responses, we observed greater than a 2-fold increase in IL-6 (all 4 animals) and IP10 (CXCL10) (2 dams, 1 infant) as reported previously (18), and the anti-inflammatory mediator IL-1RA (1 dam, 2 infants) within 1-day post SARS-CoV-2 infection (**Fig. 2B**). One infant demonstrated increased production of IL-17 and GM-CSF at several time points post-infection. Interestingly, IL-8 was upregulated greater than 7-fold in 3 of the 4 animals at day 14 post-infection. However, we did not identify any specific cytokine signature that distinguished dams from infants.

**Fig. 2.**
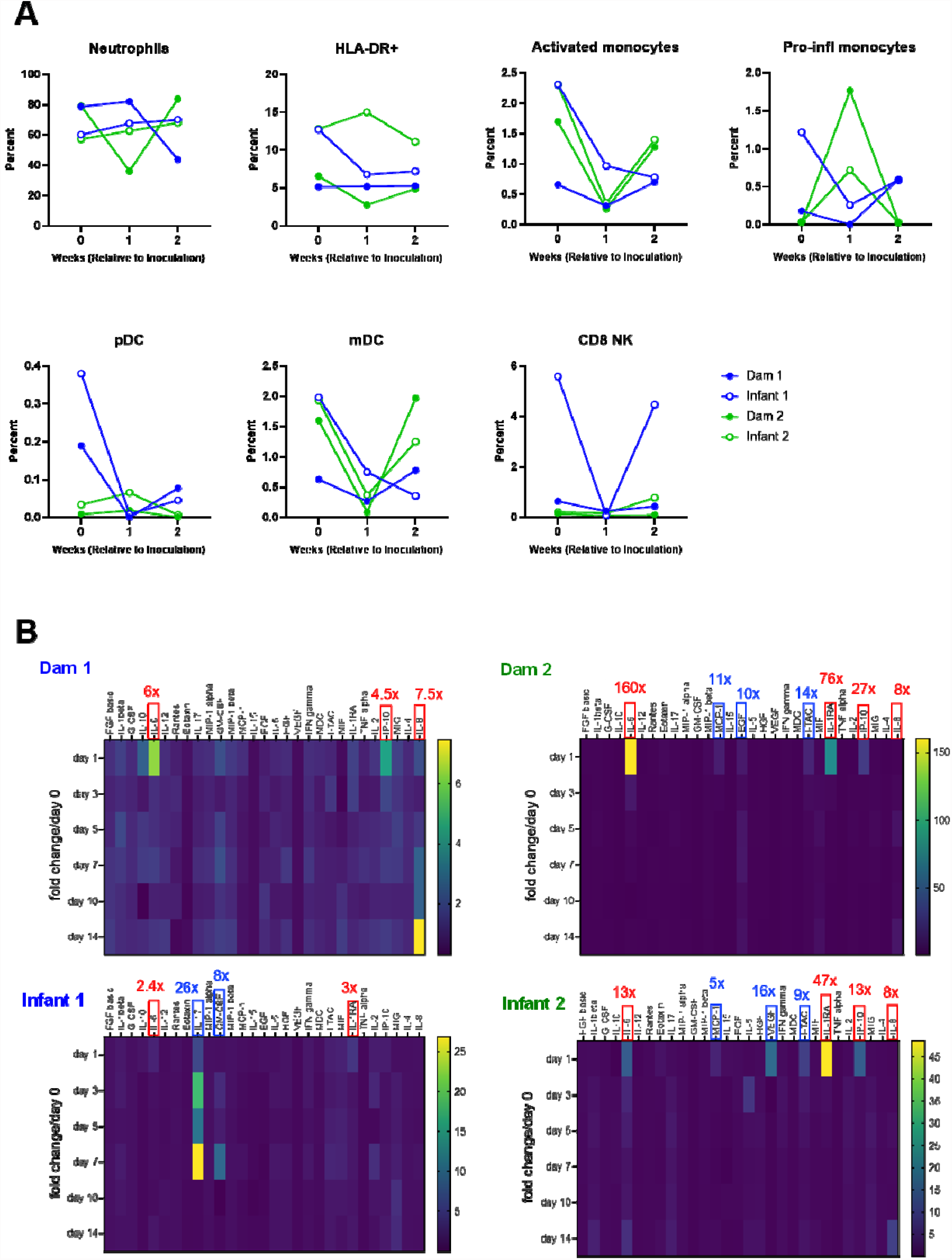
Circulating innate immune cell and cytokine responses in SARS-CoV-2-infected dam and infant rhesus macaques. **A** Kinetics of innate immune cell responses including neutrophils, total HLA-DR+ (lineage-, CD66-) cells, activated monocytes, pro-inflammatory monocytes, plasmacytoid dendritic cells (pDC), myeloid dendritic cells (mDC) and CD8^+^ natural killer (NK) cells. **B** Heat maps of innate cytokines for each animal represented as fold change over day 0 post SARS-CoV-2 infection. Notable increases in key cytokines are denoted in red box (values in red represent peak fold change over day 0). The scale to the right of the heat map is in pg/ml.

### Kinetics of neutralizing antibody (nAb) development differ between dam and infant rhesus macaques infected with SARS-CoV-2

We next measured anti-spike (S) binding and nAbs in serum and breast milk samples. With the limitation that the small group sizes preclude statistical significance, we found that while dams had higher serum S-binding IgG antibody titers than infants (**Fig. 3A**), infants demonstrated faster kinetics of nAb development. Infants had higher nAb titers at day 7 post-infection, yet similar nAb titers at day 14 post-infection compared to dams (**Fig. 3B)**. S-binding IgG, but not IgA, was detected in breast milk of infected dams at day 14 post-infection (**Fig. 3C**). NAbs in breast milk were detected in one dam at day 14 post-infection (**Fig. 3D**). A multiplexed antibody binding assay did not detect differences in receptor binding domain (RBD), S1, S2, full S and N-terminal domain (NTD)-specific antibodies between SARS-CoV-2-infected dam and infant macaques at day 14 (**Fig. 3E**).

**Fig. 3.**
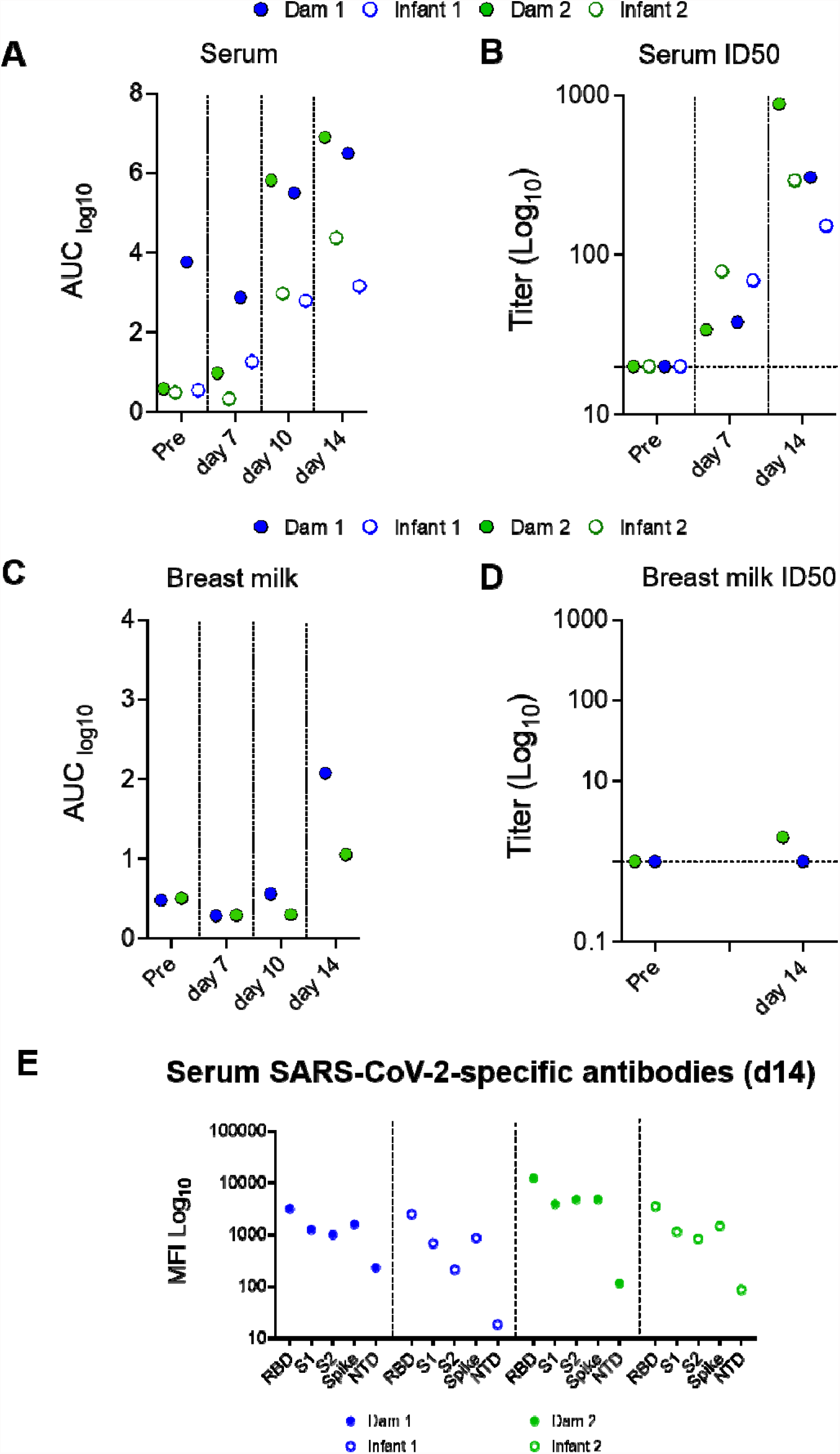
SARS-CoV-2-elicited binding and neutralizing antibody (nAb) responses in infant and dam rhesus macaques. **A** S-2P protein-specific antibody responses were measured in serum and **C** breast milk by ELISA. Serial dilutions of plasma (starting at 1:10) and breast milk (starting at 1:1) were assayed for IgG binding to SARS-CoV-2 S. Data are reported as log_10_ AUC values. **B** Neutralization capacity in serum and **D** breast milk was measured using S D614G-pseudotyped viruses and HEK 293T cells expressing ACE2 receptors. Results are expressed as reciprocal 50% inhibitory dilution (ID_50_). Gray dotted lines represent detection cutoff. **E** Antibody epitope specificity measured by binding antibody multiplex assay (BAMA). Plasma was diluted 1:1,000 to measure binding to different domains of the S protein, including the RBD, S1, S2, full-length S protein, and NTD. Binding antibody responses are reported as log_10_-transformed MFI after subtraction of background values.

### Age-dependent differences in transcriptomic responses in the trachea reveal downregulated interferon stimulated genes and cilia injury signatures in adult macaques

The hierarchical and principal component analyses and heatmap (**Supplementary Fig. 3A-C**) of the differentially expressed genes (DEGs) of tracheal cells (epithelial, immune and mesenchymal) demonstrated altered transcriptomic expression of infant and dam rhesus macaques on day 14 after SARS-CoV-2 infection, where mothers and offspring clustered with each other (**Supplementary Fig. 3C**). To identify gene ontology (GO) and hallmark gene sets associated with altered expression, gene set enrichment analysis (GSEA) was performed (1). The Hallmark pathway analysis revealed a decrease in trachea cell genes associated with the *Interferon Alpha* and *Interferon Gamma* Hallmark pathways in adult compared to infant macaques (**Supplementary Table 2A**). Specifically, *IFI6, XAF1, DDX60, OAS2, HERC6, MX2, IFI44L, IFIT1, SAMD9L, PARP14 and IFIT12* are significantly decreased in both adults compared to their infants (**Fig. 4A, Supplementary Fig. 4**).

**Fig. 4.**
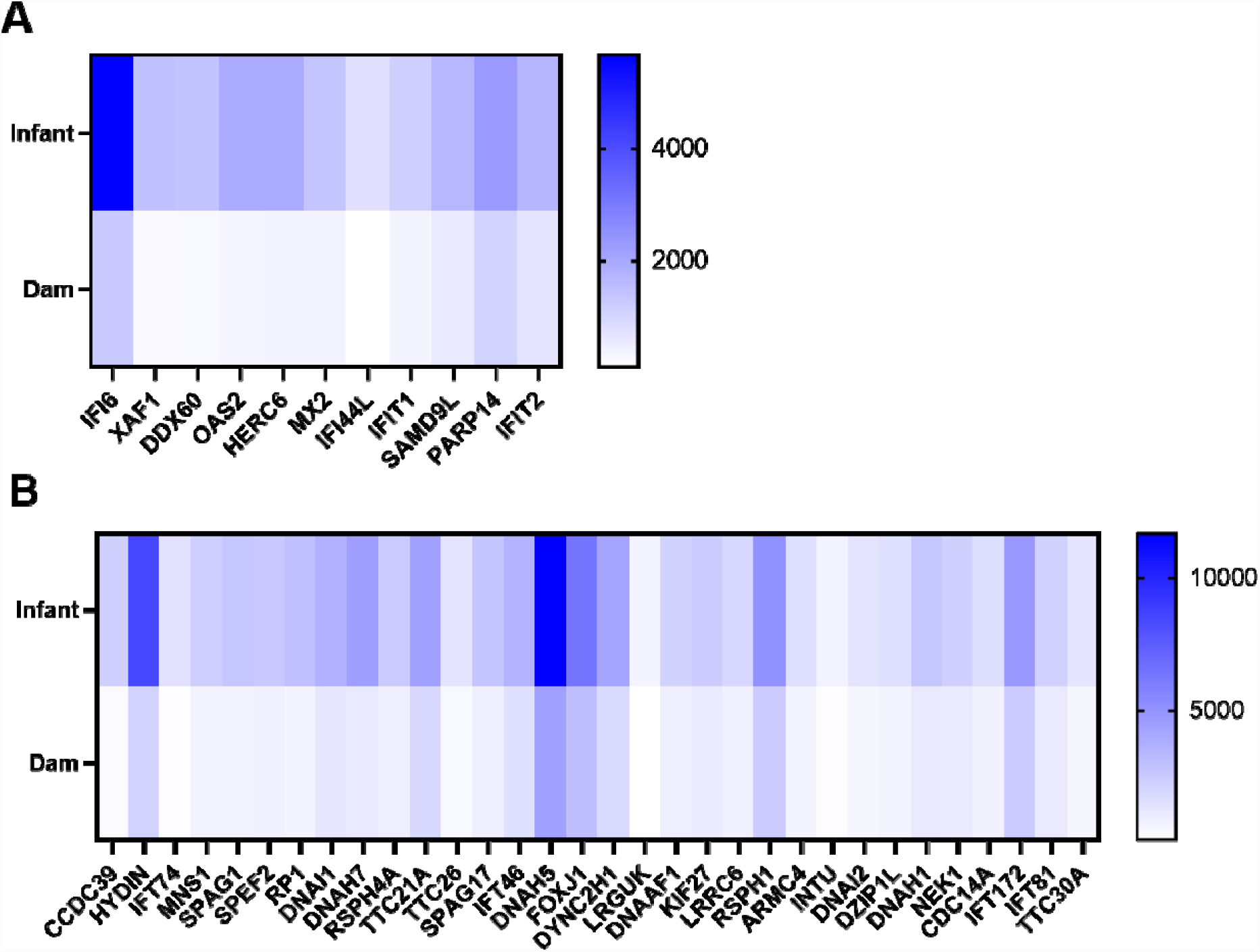
Age-dependent differences in transcriptomic responses in the trachea reveal downregulated interferon stimulated genes and cilia injury signatures in adult macaques. **A** Heat map of differentially expressed interferon stimulated genes and **B** genes related to cilia structure and function that are downregulated in adult compared with infant macaques 14 days after SARS-CoV-2 infection. Blue represents relative upregulation of gene expression and white represents relative downregulation of gene expression. Genes are arranged by log_2_ fold change with the largest log_2_ fold change to the left and the smallest log_2_ fold change to the right.

Alternatively, in the GO pathway analysis we identified an overwhelming signature of decreased cilia structure and function in adult compared to infant tracheas (**Fig. 4B, Supplementary Fig. 5**). In fact, all GO pathways that were significantly enriched among downregulated genes (FWER adjusted P-value ≤ 0.05) were related to cilia structure and motility (**Supplementary Table 2B**). Considering SARS-CoV-2 infection induces a de-differentiation of multiciliated cells (19), these results suggest a prolonged impairment of cilia functions in adult compared to infant rhesus macaques 14 days after SARS-CoV-2 infection.

### Conducting airways of SARS-CoV-2 infected adult macaques show a profibrotic transcriptomic signature

In both the GO and Hallmark pathway analyses, there was significant enrichment among upregulated genes (FWER adjusted P-value ≤ 0.05) for pathways associated with wound repair and fibrosis in adult compared with infant macaques at day 14 post infection (**Supplementary Table 3**). Of the forty GO pathways that are significantly enriched among upregulated genes in the adult macaques, the overwhelming majority were associated with extracellular matrix (ECM) organization and ECM metabolism (**Supplementary Table 3**). Similarly, the Hallmark pathway analysis revealed multiple pathways associated with wound repair and fibrosis in adult macaques including *Epithelial mesenchymal transition, Angiogenesis, Coagulation, Hypoxia, Apoptosis and TGF beta signaling* (**Supplementary Table 4**). Of the most DEGs of the upregulated pathways in adults compared to infants, all of them were associated with the ECM (**Fig. 5A, Supplementary Fig. 7**). We stained trachea tissue for protein gene product 9.5 (PGP 9.5) (also known as ubiquitin C-terminal hydrolase-L1 [UCH-L1]) (**Fig. 5D-G**) and Substance P (Sub P) (**Fig. 5 H-K**), proteins involved in wound healing (20, 21), which were both higher in infant compared to adult tracheas (**Fig. 5**).

**Fig. 5.**
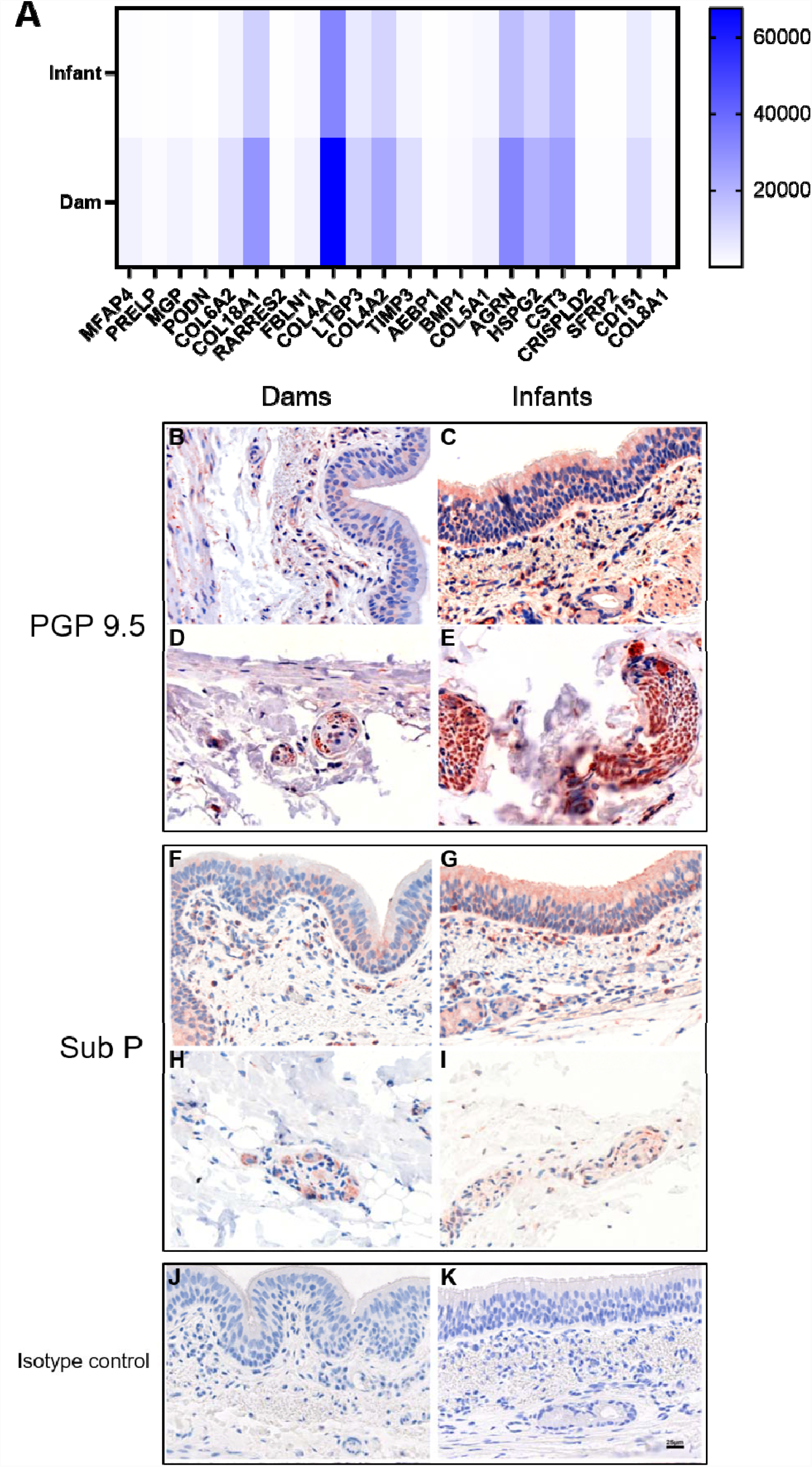
Upper airways of SARS-CoV-2 infected adult macaques have a profibrotic transcriptomic signature. **A** Heat map of differentially expressed genes related to extracellular matrix structure and metabolism that are upregulated in adult compared with infant macaques 14 days after SARS-CoV-2 infection. Blue represents relative upregulation of gene expression and white represents relative downregulation of gene expression. Genes are arranged by log_2_ fold change with the largest log_2_ fold change to the left and the smallest log_2_ fold change to the right. Assessment of protein gene product (PGP) 9.5 expression in (**B**) dam 1, (**D**) dam 2, (**E**) infant 1 and (**C**) infant 2, substance P (sub P) expression in (**F**) dam 1, (**H**) dam 2, (**I**) infant 1 and (**G**) infant 2 trachea tissues and (**J**) dam and (**K**) infant isotype controls via immunohistochemical staining.

## Discussion

In this pilot study, we evaluated age-dependent differences in SARS-CoV-2 pathogenesis by inoculating two dam-infant rhesus macaque pairs with SARS-CoV-2. We demonstrate that both infant and dam rhesus macaques became productively infected and exhibited no to mild clinical symptoms. These clinical observations are similar to those observed in other NHP studies of SARS-CoV-2 infection in adults (15, 16). Lung radiograph scores showed mild pulmonary infiltrates in both dams and infants, however only in dams were pulmonary infiltrates observed for multiple days (≥2). Histologic examination of pulmonary infiltrates was extremely minimal and largely resolving when examined day 14 post-infection. This is consistent with previous experiments (18).

While viral loads in pharyngeal swabs and BAL samples did not differ between dam and infant macaques, differences in antibody response kinetics between the dams and infants were observed. For example, while there were higher anti-spike IgG binding antibody titers in adult macaques at day 10 and 14 post-infection, infants exhibited higher nAb titers at day 7. These data suggest faster kinetics of SARS-CoV-2 nAb responses in infant macaques. In pediatric cohorts infected with SARS-CoV-2, results have been variable and levels of nAbs have been reported as lower, higher or not different compared to infected adults (22-26). However, age-dependent differences in nAb development have been observed for other pathogens. For example, in HIV-1 infected children, circulating broadly neutralizing antibodies (bnAbs) arise earlier in infection, and have higher potency and breadth compared to adults (27-32). Future work using this model should include measurement of T cell subsets in both blood and lymphoid tissue as well as determination of breadth against multiple SARS-CoV-2 variants.

Although there were not major differences in serum cytokines between adult and infant SARS-CoV-2 infected macaques, we observed increases in relevant pro-inflammatory cytokines. For example, IL-6 and IP10 (CXCL10) were increased in 3 of the 4 animals within 1-day post SARS-CoV-2, mirroring what was previously reported in rhesus macaques and other animal models of SARS-CoV-2 infection (33-35). In humans, IL-6 and IL-RA correlate with COVID-19 severity (36, 37) while CXCL10 correlates with viral load in nasopharynx (38). Interestingly, we observed a delayed increase in IL-8 in 3 of the 4 macaques at day 14 post-infection. Following SARS-CoV-2 infected macaques for greater than 14 days would be worthwhile in determining if late-onset pro-inflammatory signatures exist. Additionally, measuring cytokines in tissues would give additional insight into age-dependent local responses to infection.

There were not major differences in innate immune cell subsets between adult and infant macaques in response to SARS-CoV-2 infection. While neutrophils were higher in adult circulation prior to infection, this could be due to a stress-induced response (39). Interestingly, HLA-DR+ (lineage-, CD66+) innate immune cells were higher in infant circulation prior to infection compared to adults. Recent studies have observed differences in circulating subsets of monocytes and dendritic cells during SARS-CoV-2 infection between children and adult cohorts (13). Assessing cellular immune subsets in tissues will be an important next step to determine age-dependent cellular trafficking after SARS-CoV-2 infection.

An increase in interferon alpha and gamma transcriptomic signatures were observed in infant macaques compared to dams. There were multiple ISGs that were upregulated in both infants compared to their dams, nearly all of which were previously shown to be significantly upregulated after SARS-CoV-2 infection. Similarly, in a pediatric cohort of SARS-CoV-2-infected children, there were multiple ISGs in epithelial cells of the upper respiratory tract that were significantly increased compared to SARS-CoV-2-infected adults late during both the early (day 0-4) and late (day 5-12) phase of infection including *IRF3, TLR3, DHX58, IFIH1*, and *DDX58* (12). Considering trachea RNA-seq data was collected at day 14 post-inoculation in our study, when viral loads were not detectable anymore, it is possible that levels of some ISGs are upregulated in infants at baseline compared to adults. Indeed, this has been demonstrated in multiple pediatric cohorts where ISGs (40) and cytokines and chemokine genes (41) were increased in SARS-CoV-2-uninfected children compared to adults. Upon SARS-CoV-2 infection, children’s IFN-α responses were greater in multiple immune cell subsets including NK CD56^low^, NKT cells, neutrophils, CXCL10+ monocytes and some CD8+ T cell subsets (40). These data suggest the IFN response in children is pre-activated in epithelial cells of the upper airways and stronger in immune cell subsets compared to adults. An increase in innate immune effectors in infants and children may also be explained by ‘trained immunity’. Trained immunity results in functional reprogramming of innate immune cells which leads to heterologous protection against secondary infections (42). Considering children are infected more frequently and with a greater number of viruses, it is possible that this leads to an induction of non-specific antiviral effects (43, 44).

There was an overwhelming signature of decreased cilia structure and function-related genes in adult compared to infant macaques. It has been previously demonstrated that SARS-CoV-2 infection leads to dedifferentiation of multiciliated cells in vitro and in vivo (19). Considering trachea RNA-seq data was collected at day 14 post-inoculation in our study, a time when viral replication at mucosal sites was undetectable, it is possible that SARS-CoV-2-infected adults have delayed cilia repair. Indeed, other groups have observed delayed repair responses after pulmonary injury in adults. For example, hypoxia or LPS-induced lung injury resulted in less lung inflammation and apoptosis in neonatal compared to adult mice, which was dependent on increased NF-κβ activation (45, 46). Additionally, neonatal mice had reduced lung permeability and pulmonary barrier disruption compared to adult mice after LPS administration, despite similar degrees of inflammation and cellular apoptosis (47). These data and our results support the hypothesis that the impacts of SARS-CoV-2 infection in the airways, including inflammation, apoptosis and barrier permeability (that implicates cilia structure and function), are lessened in neonates and young children compared to adults.

We observed that multiple genes and pathways associated with ECM composition and metabolism, coagulation, angiogenesis and hypoxia are increased in the trachea of dams compared to infants. These profibrotic transcriptomic signatures suggest greater activation of the cellular repair process in response to injury and apoptosis in the upper airway in adults compared to infants. Indeed, the deposition of multiple components of the lung ECM including hyaluronan and fibrinogen (48-50) as well as profibrotic macrophage accumulation (50) has been associated with severe COVID-19 in adults. Additionally, an increase in ECM-remodeling enzymes is associated with lethal immunopathology during severe influenza infection (51). Alterations in ECM can be not only a consequence of lung fibrosis but also a driver of its progression (52) and adult acute respiratory distress syndrome (ARDS) is more often associated with permanent alveolar simplification and fibrosis compared to infants or children (53). This likely explains why children with ARDS experience less morbidity and mortality outcomes compared to adults (54). An increase in ECM and collagen genes in the adult macaques may be indicative of a dysregulation of injury repair. Targeting the ECM may be a therapeutic strategy for promoting injury repair in adults with severe COVID-19.

Our study has limitations. Firstly, we only infected two dam-infant macaque pairs and more animal numbers are needed to determine statistical differences. Additionally, a mock-inoculated control group is necessary to decipher whether the differences observed are due to SARS-CoV-2 infection alone, the age-dependent maturation of tissues and the immune response, and/or experimental procedures. Finally, the time of euthanasia was not focused on evaluating acute inflammatory responses in tissues, as at 2 weeks, virus replication is mainly gone and tissue responses reflect repair in this animal model, and we evaluated trachea instead of lung responses. However, our study is valuable in that it agrees with currently published data in SARS-CoV-2-infected pediatric and adult cohorts and furthers our understanding of why younger populations are less susceptible to severe COVID-19 compared to adults. Additionally, this model will allow further definition of molecular mechanisms of age-dependent SARS-CoV-2 pathogenesis and assess efficacy of medical countermeasures.

## Methods

### Animals, SARS-CoV-2 infection and sample collection

The four Indian origin rhesus macaques (Macaca mulatta) used in this study were housed at CNRPC in accordance with the recommendations of the Association for Assessment and Accreditation of Laboratory Animal Care International Standards and with the recommendations in the Guide for the Care and Use of Laboratory Animals of the United States - National Institutes of Health. The Institutional Animal Use and Care Committee approved these experiments (study protocol# 21702). All animals were challenged through combined intratracheal (IT, 2.0 mL for dams, 1 ml for infants) and intranasal (IN, 0.2 5 mL per nostril) inoculation with an infectious dose of 2.5 × 10^6 PFU for the dams and 1.5 × 10^6 PFU for the infants of SARS-CoV-2 (2019-nCoV/USA-WA1/2020). The stock was obtained from BEI Resources (NR-52281). The stock underwent deep sequencing to confirm homology with the WA1/2020 isolate. Virus was stored at -80°C prior to use, thawed rapidly at 37°C, and placed immediately on wet ice. Nasal swabs, pharyngeal swabs, BAL, blood, breast milk and rectal swab samples were collected 1 week before infection and every 2-3 days post-challenge. Nasopharyngeal, oropharyngeal and buccal secretions were collected with FLOQSwabs^™^ (Copan), placed in a vial with DNA/RNA Shield^™^ solution (Zymo Research), and stored at -70°C until further processing.

BAL was performed using a 20F rubber feeding tube with instillation of 20 ml sterile physiologic saline followed by aspiration with a syringe. BAL samples were spun in the lab. The BAL cell pellet, together with 0.5 ml of supernatant, was then mixed with 1.5 ml of TRIzol®-LS (Thermo Fisher Scientific) and cryopreserved at -70° C. Additional aliquots of BAL supernatant were also immediately cryopreserved. Blood and milk were collected and processed as previously described (55-57).

Radiographs were obtained with a HF100+ Ultralight imaging unit (MinXRay, Northbrook, IL) at 50 kVp, 40mA, and 0.1 sec. Ventrodorsal, dorsoventral, R lateral, and L lateral radiographs were obtained prior to inoculation and on days 1, 3, 5, 7, 10, and 14 post inoculation. Radiographs were scored for the presence of pulmonary infiltrates by a board-certified veterinary radiologist, who was blinded to the experimental group and time point, according to a standard scoring system (0: normal; 1: mild interstitial pulmonary infiltrates; 2: moderate pulmonary infiltrates perhaps with partial cardiac border effacement and small areas of pulmonary consolidation; 3: severe interstitial infiltrates, large areas of pulmonary consolidation, alveolar patterns and air bronchograms). Individual lobes were scored and scores per animal per day were totaled.

At the end of the study, animals were euthanized, and a full necropsy was performed for tissue collection, including trachea for cell isolation and fixed lung tissues for histopathology.

### Trachea collection and cell processing

Upon necropsy, tracheobronchial tissues were collected. Airway epithelial cells were isolated by placing tracheas in Minimum Essential Medium Eagle; Joklik Modification (Lonza) containing 0.1% type XIV protease (Sigma-Aldrich), 50 U/mL penicillin, 50 μg/mL each streptomycin and gentamicin, and 100 µg/mL geneticin G418 (Invitrogen) overnight. Cells were gently aspirated off and stored in liquid nitrogen prior to RNA isolation.

### Viral load assay

SARS-CoV-2 infection was determined by quantitative Polymerase Chain Reaction (qPCR) of SARS-CoV-2 Genomic (orf1a) and Subgenomic (N) RNA.

#### SARS-CoV-2 Genomic (orf1a) qPCR

A QIAsymphony SP (Qiagen, Hilden, Germany) automated sample preparation platform along with a virus/pathogen DSP midi kit and the complex800 protocol were used to extract viral RNA from 800 µL of respiratory sample. A reverse primer specific to the orf1a sequence of SARS-CoV-2 (5’-CGTGCCTACAGTACTCAGAATC-3’) was annealed to the extracted RNA and then reverse transcribed into cDNA using SuperScript™ III Reverse Transcriptase (Thermo Fisher Scientific, Waltham, MA) along with RNAse Out (Thermo Fisher Scientific, Waltham, MA). The resulting cDNA was then treated with Rnase H (Thermo Fisher Scientific, Waltham, MA) and added to a custom 4x TaqMan™ Gene Expression Master Mix (Thermo Fisher Scientific, Waltham, MA) containing primers and a fluorescently labeled hydrolysis probe specific for the orf1a sequence of SARS-CoV-2 (forward primer 5’-GTGCTCATGGATGGCTCTATTA-3’, reverse primer 5’-CGTGCCTACAGTACTCAGAATC-3’, probe 5’-/56-FAM/ ACCTACCTT/ZEN/GAAGGTTCTGTTAGAGTG GT/3IABkFQ/-3). All PCR setup steps were performed using QIAgility instruments (Qiagen, Hilden, Germany). The qPCR was then carried out on a QuantStudio 3 Real-Time PCR System (Thermo Fisher Scientific, Waltham, MA). SARS-CoV-2 genomic (orf1a) RNA copies per reaction were interpolated using quantification cycle data and a serial dilution of a highly characterized custom RNA transcript containing the SARS-CoV-2 orf1a sequence. Mean RNA copies per milliliter were then calculated by applying the assay dilution factor (DF=11.7). The limit of quantification (LOQ) for this assay is approximately 31 RNA cp/mL (1.49 log10) with 800 uL of sample.

#### Subgenomic (N) RNA qPCR assay

A QIAsymphony SP (Qiagen, Hilden, Germany) automated sample preparation platform along with a virus/pathogen DSP midi kit and the complex800 protocol were used to extract viral RNA from 800 µL of respiratory sample. The extracted RNA was then added to TaqMan™ Fast Virus 1-Step Master Mix (Thermo Fisher Scientific, Waltham, MA) containing primers and a fluorescently labeled hydrolysis probe specific for mRNA from the nucleocapsid gene of SARS-CoV-2 (forward primer 5’-CGATCTCTTGTAGATCTGTTCTC-3’, reverse primer 5’-GGTGAACCAAGACGCAGTAT-3’, probe 5’-/56-FAM/TAACCAGAA/ZEN/TGGAGAACGCAGT GGG/3IABkFQ/-3’). All PCR setup steps were performed using QIAgility instruments (Qiagen, Hilden, Germany). The qPCR was carried out on a QuantStudio 3 Real-Time PCR System (Thermo Fisher Scientific, Waltham, MA). SARS-CoV-2 subgenomic (N) RNA copies per reaction were interpolated using quantification cycle data and a serial dilution of a highly characterized custom RNA transcript containing the SARS-CoV-2 subgenomic nucleocapsid sequence. Mean RNA copies per milliliter were then calculated by applying the assay dilution factor (DF=5.625). The limit of quantification (LOQ) for this assay is approximately 31 RNA cp/mL (1.49 log10) with 800 uL of sample.

### Spike-specific IgG ELISA

IgG binding to the stabilized SARS-CoV-2 S protein S-2P was measured in plasma using ELISA as previously described (58). Three hundred eighty-four–well plates were coated overnight with S protein (2 μg/ml) produced by the Protein Production Facility (PPF) at the Duke Human Vaccine Institute. Plates were then blocked with assay diluent (PBS containing 4% whey, 15% normal goat serum, and 0.5% Tween 20). Ten serial fourfold dilutions starting at 1:10 for plasma and undiluted for breast milk were added to the plates and incubated for 1 hour, followed by detection with a horseradish peroxidase (HRP)–conjugated mouse anti-monkey IgG (SouthernBiotech). The plates were developed by using an 2,2′-azinobis(3-ethylbenzthiazolinesulfonic acid peroxidase substrate system (Colonial Scientific), and absorbance was read at 450 nm with a SpectraMax microplate reader (Molecular Devices). Results were calculated as AUC and EC_50_ values. AUC values were calculated using the trapezoidal rule. EC_50_ values were calculated by fitting a four-parameter logistic function using nonlinear regression. Pooled NHP convalescent serum to SARS-CoV-2 (BEI Resources, NR-52401) was used in all IgG assays to ensure interassay reproducibility, but standard curves were not developed given the lack of an SARS-CoV-2 RM-specific IgG reagent of known concentration. IgA binding to S2P was measured following a similar ELISA protocol but using as detection Ab anti-rhesus IgA 10F12-biotin (NHP Reagent Resource catalog AB_2819304) followed by Streptadividin-HRP (Pierce, catalog 21126).

### Binding antibody multiplex assay

SARS-CoV-2 antigens, including whole S (produced by PPF), S1 (Sino Biological, catalog no. 40591-V08H), S2 (Sino Biological, catalog no. 40590-V08B), RBD (Sino Biological, catalog no. 40592-V08H), and NTD (Sino Biological, catalog no. 40591-V49H) were conjugated to Magplex beads (Bio-Rad, Hercules, CA). The conjugated beads were incubated on filter plates (Millipore, Stafford, VA) for 30 min before plasma samples were added. Plasma samples were diluted in assay diluent [1% dry milk, 5% goat serum, and 0.05% Tween 20 in PBS (pH 7.4.)] at a 1:1000-point dilution. Beads and diluted samples were incubated for 30 min with gentle rotation, and IgG binding was detected using a PE-conjugated mouse anti-monkey IgG (SouthernBiotech, Birmingham, Alabama) at 2 µg/ml. Plates were washed and acquired on a Bio-Plex 200 instrument (Bio-Rad, Hercules, CA), and IgG binding was reported as mean fluorescence intensity (MFI). To assess assay background, the MFIs of wells without sample (blank wells) were used, and nonspecific binding of the samples to unconjugated blank beads was evaluated.

### Pseudovirus Ab neutralization assay

SARS-CoV-2 neutralization was assessed with S-pseudotyped viruses in 293 T/ACE2 cells as a function of reductions in luciferase (Luc) reporter activity. 293 T/ACE2 cells were provided by M. Farzan and H. Mu at Scripps Florida. Cells were maintained in Dulbecco’s modified Eagle’s medium containing 10% fetal bovine serum (FBS), 25 mM Hepes, gentamycin (50 μg/ml), and puromycin (3 μg/ml). An expression plasmid encoding codon-optimized full-length S of the Wuhan-1 strain (VRC7480) was provided by B.S.G. and K.S.C. at the Vaccine Research Center, NIH (USA). The D614G amino acid change was introduced into VRC7480 by site-directed mutagenesis using the QuikChange Lightning Site-Directed Mutagenesis Kit from Agilent Technologies (catalog no. 210518). The mutation was confirmed by full-length S gene sequencing. Pseudovirions were produced in HEK293 T/17 cells (American Type Culture Collection, catalog no. CRL-11268) by transfection using Fugene 6 (Promega, catalog no. E2692) and a combination of S plasmid, lentiviral backbone plasmid (pCMV ΔR8.2), and firefly Luc reporter gene plasmid (pHR’ CMV Luc) (78) in a 1:17:17 ratio. Transfections were allowed to proceed for 16 to 20 hours at 37°C. Medium was removed, monolayers were rinsed with growth medium, and 15 ml of fresh growth medium was added. Pseudovirus-containing culture medium was collected after an additional 2 days of incubation and was clarified of cells by low-speed centrifugation and 0.45-μm filtration and stored in aliquots at −80°C. Median tissue culture infectious dose assays were performed on thawed aliquots to determine the infectious dose for neutralization assays.

For neutralization, a pretitrated dose of pseudovirus was incubated with eight serial fivefold dilutions of serum samples in duplicate in a total volume of 150 μl for 1 hour at 37°C in 96-well flat-bottom poly-L-lysine–coated culture plates (Corning Biocoat). HEK 293T cells expressing ACE2 receptors were suspended using TrypLE Select Enzyme solution (Thermo Fisher Scientific) and immediately added to all wells (10,000 cells in 100 μl of growth medium per well). One set of eight control wells received cells and virus (virus control), and another set of eight wells received cells only (background control). After 66 to 72 hours of incubation, medium was removed by gentle aspiration and 30 μl of Promega 1X lysis buffer was added to all wells. After a 10-min incubation at RT, 100 μl of Bright-Glo Luc reagent was added to all wells. After 1 to 2 min, 110 μl of the cell lysate was transferred to a black/white plate (PerkinElmer). Luminescence was measured using a PerkinElmer Life Sciences, Model Victor2 luminometer. Neutralization titers are the serum dilution at which relative light units (RLUs) were reduced by either 50% (ID_50_) or 80% (ID_80_) compared with virus control wells after subtraction of background RLUs. Serum samples were heat-inactivated for 30 min at 56°C before assay.

### RNA isolation and transcript analysis

Tracheal cells (epithelial, immune and mesenchymal) were lysed in 350 ul of Trizol, and RNA was extracted by phenol:chloroform method, and collected over RNAeasy column (Qiagen, Germantown, MD). The RNA concentration and integrity was determined on the NanoDrop ND-1000 (Thermo Fisher Scientific). 1000ng RNA from each sample was used as a template for preparing Illumina compatible libraries using the TruSeq RNA Library Prep Kit v2 (Illumina). Library sizes were checked using D5000 high sensitivity tape on the TapesSation 2200 (Agilent), and pooled libraries concentration were determined by Qubit 3.0 Fluorometer (Thermo Fisher Scientific). A library input of 1.8 pM with 1% PhiX (Illumina) spike-in was sequenced using the NextSeq 500 instrument with the NextSeq500/550 High Output v2.5 Kit (Illumina).

### Analysis of the bulk RNA-seq data

RNA-Seq data was quality checked with FastQC (59) and preprocessing was carried out using TrimGalore (60) toolkit to trim low-quality bases and Illumina adapter sequences using default settings. Reads were aligned to the ENSEMBL Homo_sapiens.GRCh38.dna.primary_assembly genome using the ENSEMBL Homo_sapiens.GRCh38.100 transcript (61) annotation file with STAR (62) splice-aware RNA-seq alignment tool in paired mode allowing maximum multimapping of 3. Gene level counts were quantified using FeatureCounts (63) tool, counting unique features in non-stranded mode and retaining both gene ID, gene name, and gene biotype mapping from the ENSEMBL annotation file. Prior to differential expression analysis, count data was collapsed to donor level and genes for which mean raw count was at least 15 were kept. Normalization and differential expression were carried out with DESeq2 (64) Bioconductor (65) package, utilizing the ‘apeglm’ Bioconductor package (66) for log fold change shrinkage, in R statistical programming environment. The design formula was constructed to test for the effect of treatment while controlling for donor.

The RNA-seq data was processed using the TrimGalore toolkit^1^ which employs Cutadapt^2^ to trim low quality bases and Illumina sequencing adapters from the 3’ end of the reads. Only reads that were 20nt or longer after trimming were kept for further analysis. Reads were mapped to the GRCh38v93 version of the human genome and transcriptome^3^ using the STAR RNA-seq alignment tool^4^. Reads were kept for subsequent analysis if they mapped to a single genomic location. Gene counts were compiled using the HTSeq tool^5^. Only genes that had at least 10 reads in any given library were used in subsequent analysis. Normalization and differential expression was carried out using the DESeq2^6^ Bioconductor^7^ package with the R statistical programming environment^8^. The false discovery rate was calculated to control for multiple hypothesis testing. Gene set enrichment analysis^9^ was performed to identify gene ontology terms and pathways associated with altered gene expression for each of the comparisons performed.

### Histopathology

The lungs were harvested and each lobe separated. All lobes were cannulated with 18-gauge blunt needles. All lobes were slowly infused with neutral buffered formalin at 30 cm fluid pressure. Once fully inflated (approx. 30 mins) the main bronchus was tied off and the lungs were placed in individual jars of formalin and fixed for 72 hours. Then they were sliced from the hilus towards the periphery into slabs approximately 5mm thick. Each slab was placed into a cassette recording its position in the stack and with further division of the slab into smaller pieces if required to fit into the cassette. Tissues were then held in 70% ethanol until processing and paraffin embedding followed by sectioning at 5 μm and generation of hematoxylin and eosin (H&E) and Masson trichrome stained slides. Slides from every other slab of the right and left caudal lobes were examined independently by 2 ACVP board certified pathologists in a blinded manner.

### Immunohistochemistry (IHC)

Substance P (Thermo Fisher) and protein gene product 9.5 (Millipore Sigma) antibodies were applied to 5µm paraffin sections. The EnVision system (Agilent) was used as the detection system with AEC (Agilent) as a chromogen. Paraffin sections were treated in an antigen unmasking solution (Vector) at 100°C for 20 minutes before incubation with a primary antibody. The slides were counterstained with Gill’s hematoxylin (StatLab). Primary antibodies were replaced by rabbit isotype control (Thermo Fisher) and run with each staining series as the negative controls.

### Biocontainment and biosafety

All work described here was performed with approved standard operating procedures for SARS-CoV-2 in a biosafety level 3 (BSL-3) facility conforming to requirements recommended in the Microbiological and Biomedical Laboratories, by the U.S. Department of Health and Human Service, the U.S. Public Health Service, and the U.S. Center for Disease Control and Prevention (CDC), and the National Institutes of Health (NIH).

## Author contributions

S.N.L., C.G., C.P., T.T., T.D., Z.M., R.R., K.J.O., R.L.S., Y.S.L., J.W.R., J.W., J.U., R.I. and R.P. performed experiments and analyzed the data, S.N.L. wrote the initial draft of the manuscript, S.N.L., S.S.I., S.R.P., L.A.M., K.K.A.V.R. contributed to study design, data analysis and editing of the manuscript. M.B. oversaw the planning and direction of the project including analysis and interpretation of the data and editing of the manuscript. All authors edited the manuscript and approved its final version.

## Acknowledgements

The work was supported by a CNPRC Pilot Program award, the Office of Research Infrastructure Program, Office of The Director, National Institutes of Health under Award Number P51OD011107, and the Duke Precision Genomics Collaboratory COVID-19 Early Career Investigator Pilot Grant Award.

## References

1. Roser MRH. Coronavirus Disease (COVID-19). https://ourworldindata.org/coronavirus:.

2. Wu Z, and McGoogan JM. Characteristics of and Important Lessons From the Coronavirus Disease 2019 (COVID-19) Outbreak in China: Summary of a Report of 72lJ314 Cases From the Chinese Center for Disease Control and Prevention. Jama. 2020;323(13):1239–42.

3. Zhu N, Zhang D, Wang W, Li X, Yang B, Song J, et al. A Novel Coronavirus from Patients with Pneumonia in China, 2019. N Engl J Med. 2020;382(8):727–33.

4. Holshue ML, DeBolt C, Lindquist S, Lofy KH, Wiesman J, Bruce H, et al. First Case of 2019 Novel Coronavirus in the United States. N Engl J Med. 2020;382(10):929–36.

5. Denison MR. Severe acute respiratory syndrome coronavirus pathogenesis, disease and vaccines: an update. Pediatr Infect Dis J. 2004;23(11 Suppl):S207–14.

6. Assiri A, McGeer A, Perl TM, Price CS, Al Rabeeah AA, Cummings DA, et al. Hospital outbreak of Middle East respiratory syndrome coronavirus. N Engl J Med. 2013;369(5):407–16.

7. Khuri-Bulos N, Payne DC, Lu X, Erdman D, Wang L, Faouri S, et al. Middle East respiratory syndrome coronavirus not detected in children hospitalized with acute respiratory illness in Amman, Jordan, March 2010 to September 2012. Clin Microbiol Infect. 2014;20(7):678–82.

8. Donnelly CA, Ghani AC, Leung GM, Hedley AJ, Fraser C, Riley S, et al. Epidemiological determinants of spread of causal agent of severe acute respiratory syndrome in Hong Kong. Lancet. 2003;361(9371):1761–6.

9. Yang X, Yu Y, Xu J, Shu H, Xia J, Liu H, et al. Clinical course and outcomes of critically ill patients with SARS-CoV-2 pneumonia in Wuhan, China: a single-centered, retrospective, observational study. Lancet Respir Med. 2020.

10. Sheikh A, McMenamin J, Taylor B, Robertson C, Public Health S, and the EIIC. SARS-CoV-2 Delta VOC in Scotland: demographics, risk of hospital admission, and vaccine effectiveness. Lancet. 2021;397(10293):2461–2.

11. Chou J, Thomas PG, and Randolph AG. Immunology of SARS-CoV-2 infection in children. Nature immunology. 2022;23(2):177–85.

12. Loske J, Rohmel J, Lukassen S, Stricker S, Magalhaes VG, Liebig J, et al. Pre-activated antiviral innate immunity in the upper airways controls early SARS-CoV-2 infection in children. Nat Biotechnol. 2021.

13. Neeland MR, Bannister S, Clifford V, Dohle K, Mulholland K, Sutton P, et al. Innate cell profiles during the acute and convalescent phase of SARS-CoV-2 infection in children. Nat Commun. 2021;12(1):1084.

14. Sposito B, Broggi A, Pandolfi L, Crotta S, Clementi N, Ferrarese R, et al. The interferon landscape along the respiratory tract impacts the severity of COVID-19. Cell. 2021;184(19):4953–68 e16.

15. Chandrashekar A, Liu J, Martinot AJ, McMahan K, Mercado NB, Peter L, et al. SARS-CoV-2 infection protects against rechallenge in rhesus macaques. Science. 2020.

16. Williamson BN, Feldmann F, Schwarz B, Meade-White K, Porter DP, Schulz J, et al. Clinical benefit of remdesivir in rhesus macaques infected with SARS-CoV-2. Nature. 2020.

17. Van Rompay KKA, Olstad KJ, Sammak RL, Dutra J, Watanabe JK, Usachenko JL, et al. Early treatment with a combination of two potent neutralizing antibodies improves clinical outcomes and reduces virus replication and lung inflammation in SARS-CoV-2 infected macaques. PLoS pathogens. 2021;17(7):e1009688.

18. Shaan Lakshmanappa Y, Elizaldi SR, Roh JW, Schmidt BA, Carroll TD, Weaver KD, et al. SARS-CoV-2 induces robust germinal center CD4 T follicular helper cell responses in rhesus macaques. Nat Commun. 2021;12(1):541.

19. Robinot R, Hubert M, de Melo GD, Lazarini F, Bruel T, Smith N, et al. SARS-CoV-2 infection induces the dedifferentiation of multiciliated cells and impairs mucociliary clearance. Nat Commun. 2021;12(1):4354.

20. Leal EC, Carvalho E, Tellechea A, Kafanas A, Tecilazich F, Kearney C, et al. Substance P promotes wound healing in diabetes by modulating inflammation and macrophage phenotype. The American journal of pathology. 2015;185(6):1638–48.

21. Liu H, Povysheva N, Rose ME, Mi Z, Banton JS, Li W, et al. Role of UCHL1 in axonal injury and functional recovery after cerebral ischemia. 2019;116(10):4643–50.

22. Weisberg SP, Connors TJ, Zhu Y, Baldwin MR, Lin WH, Wontakal S, et al. Distinct antibody responses to SARS-CoV-2 in children and adults across the COVID-19 clinical spectrum. Nature immunology. 2021;22(1):25–31.

23. Garrido C, Hurst JH, Lorang CG, Aquino JN, Rodriguez J, Pfeiffer TS, et al. Asymptomatic or mild symptomatic SARS-CoV-2 infection elicits durable neutralizing antibody responses in children and adolescents. JCI insight. 2021;6(17).

24. Yang HS, Costa V, Racine-Brzostek SE, Acker KP, Yee J, Chen Z, et al. Association of Age With SARS-CoV-2 Antibody Response. JAMA network open. 2021;4(3):e214302.

25. Pierce CA, Preston-Hurlburt P, Dai Y, Aschner CB, Cheshenko N, Galen B, et al. Immune responses to SARS-CoV-2 infection in hospitalized pediatric and adult patients. Sci Transl Med. 2020;12(564):eabd5487.

26. Dowell AC, Butler MS, Jinks E, Tut G, Lancaster T, Sylla P, et al. Children develop robust and sustained cross-reactive spike-specific immune responses to SARS-CoV-2 infection. Nature immunology. 2022;23(1):40–9.

27. Muenchhoff M, Adland E, Karimanzira O, Crowther C, Pace M, Csala A, et al. Nonprogressing HIV-infected children share fundamental immunological features of nonpathogenic SIV infection. Sci Transl Med. 2016;8(358):358ra125.

28. Ditse Z, Muenchhoff M, Adland E, Jooste P, Goulder P, Moore PL, et al. HIV-1 Subtype C-Infected Children with Exceptional Neutralization Breadth Exhibit Polyclonal Responses Targeting Known Epitopes. Journal of virology. 2018;92(17).

29. Goo L, Chohan V, Nduati R, and Overbaugh J. Early development of broadly neutralizing antibodies in HIV-1-infected infants. Nature medicine. 2014;20(6):655–8.

30. Makhdoomi MA, Khan L, Kumar S, Aggarwal H, Singh R, Lodha R, et al. Evolution of cross-neutralizing antibodies and mapping epitope specificity in plasma of chronic HIV-1-infected antiretroviral therapy-naïve children from India. The Journal of general virology. 2017;98(7):1879–91.

31. Mishra N, Sharma S, Dobhal A, Kumar S, Chawla H, Singh R, et al. Broadly neutralizing plasma antibodies effective against autologous circulating viruses in infants with multivariant HIV-1 infection. Nat Commun. 2020;11(1):4409.

32. Kumar S, Panda H, Makhdoomi MA, Mishra N, Safdari HA, Chawla H, et al. An HIV-1 Broadly Neutralizing Antibody from a Clade C-Infected Pediatric Elite Neutralizer Potently Neutralizes the Contemporaneous and Autologous Evolving Viruses. Journal of virology. 2019;93(4).

33. Francis ME, Goncin U, Kroeker A, Swan C, Ralph R, Lu Y, et al. SARS-CoV-2 infection in the Syrian hamster model causes inflammation as well as type I interferon dysregulation in both respiratory and non-respiratory tissues including the heart and kidney. PLoS pathogens. 2021;17(7):e1009705.

34. Winkler ES, Bailey AL, Kafai NM, Nair S, McCune BT, Yu J, et al. SARS-CoV-2 infection of human ACE2-transgenic mice causes severe lung inflammation and impaired function. Nature immunology. 2020;21(11):1327–35.

35. Munster VJ, Feldmann F, Williamson BN, van Doremalen N, Pérez-Pérez L, Schulz J, et al. Respiratory disease in rhesus macaques inoculated with SARS-CoV-2. Nature. 2020;585(7824):268–72.

36. Del Valle DM, Kim-Schulze S, Huang H-H, Beckmann ND, Nirenberg S, Wang B, et al. An inflammatory cytokine signature predicts COVID-19 severity and survival. Nature medicine. 2020;26(10):1636–43.

37. Zhao Y, Qin L, Zhang P, Li K, Liang L, Sun J, et al. Longitudinal COVID-19 profiling associates IL-1RA and IL-10 with disease severity and RANTES with mild disease. JCI insight. 2020;5(13).

38. Cheemarla NR, Watkins TA, Mihaylova VT, Wang B, Zhao D, Wang G, et al. Dynamic innate immune response determines susceptibility to SARS-CoV-2 infection and early replication kinetics. The Journal of experimental medicine. 2021;218(8).

39. Ince LM, Weber J, and Scheiermann C. Control of Leukocyte Trafficking by Stress-Associated Hormones. 2019;9.

40. Yoshida M, Worlock KB, Huang N, Lindeboom RGH, Butler CR, Kumasaka N, et al. Local and systemic responses to SARS-CoV-2 infection in children and adults. Nature. 2021.

41. Loske J, Röhmel J, Lukassen S, Stricker S, Magalhães VG, Liebig J, et al. Pre-activated antiviral innate immunity in the upper airways controls early SARS-CoV-2 infection in children. Nature Biotechnology. 2021.

42. Netea MG, Domínguez-Andrés J, Barreiro LB, Chavakis T, Divangahi M, Fuchs E, et al. Defining trained immunity and its role in health and disease. Nature Reviews Immunology. 2020;20(6):375–88.

43. Elahi S. Neonatal and Children’s Immune System and COVID-19: Biased Immune Tolerance versus Resistance Strategy. 2020;205(8):1990–7.

44. Midulla F, Cristiani L, and Mancino E. Will children reveal their secret? The coronavirus dilemma. The European respiratory journal. 2020;55(6).

45. Alvira CM, Abate A, Yang G, Dennery PA, and Rabinovitch M. Nuclear factor-kappaB activation in neonatal mouse lung protects against lipopolysaccharide-induced inflammation. American journal of respiratory and critical care medicine. 2007;175(8):805–15.

46. Yang G, Abate A, George AG, Weng YH, and Dennery PA. Maturational differences in lung NF-kappaB activation and their role in tolerance to hyperoxia. The Journal of clinical investigation. 2004;114(5):669–78.

47. Ying L, Alvira CM, and Cornfield DN. Developmental differences in focal adhesion kinase expression modulate pulmonary endothelial barrier function in response to inflammation. American journal of physiology Lung cellular and molecular physiology. 2018;315(1):L66–l77.

48. Hellman U, Karlsson MG, Engström-Laurent A, Cajander S, Dorofte L, Ahlm C, et al. Presence of hyaluronan in lung alveoli in severe Covid-19: An opening for new treatment options? The Journal of biological chemistry. 2020;295(45):15418–22.

49. Wool GD, and Miller JL. The Impact of COVID-19 Disease on Platelets and Coagulation. Pathobiology : journal of immunopathology, molecular and cellular biology. 2021;88(1):15–27.

50. Wendisch D, Dietrich O, Mari T, von Stillfried S, Ibarra IL, Mittermaier M, et al. SARS-CoV-2 infection triggers profibrotic macrophage responses and lung fibrosis. Cell. 2021;184(26):6243–61.e27.

51. Boyd DF, Allen EK, Randolph AG, Guo X-zJ, Weng Y, Sanders CJ, et al. Exuberant fibroblast activity compromises lung function via ADAMTS4. Nature. 2020;587(7834):466–71.

52. Herrera J, Henke CA, and Bitterman PB. Extracellular matrix as a driver of progressive fibrosis. The Journal of clinical investigation. 2018;128(1):45–53.

53. Im D, Shi W, and Driscoll B. Pediatric Acute Respiratory Distress Syndrome: Fibrosis versus Repair. Front Pediatr. 2016;4:28-.

54. Orloff KE, Turner DA, and Rehder KJ. The Current State of Pediatric Acute Respiratory Distress Syndrome. Pediatric allergy, immunology, and pulmonology. 2019;32(2):35–44.

55. Permar SR, Wilks AB, Ehlinger EP, Kang HH, Mahlokozera T, Coffey RT, et al. Limited contribution of mucosal IgA to Simian immunodeficiency virus (SIV)-specific neutralizing antibody response and virus envelope evolution in breast milk of SIV-infected, lactating rhesus monkeys. Journal of virology. 2010;84(16):8209–18.

56. Jensen K, Nabi R, Van Rompay KKA, Robichaux S, Lifson JD, Piatak M, Jr., et al. Vaccine-Elicited Mucosal and Systemic Antibody Responses Are Associated with Reduced Simian Immunodeficiency Viremia in Infant Rhesus Macaques. Journal of virology. 2016;90(16):7285–302.

57. Marthas ML, Van Rompay KK, Abbott Z, Earl P, Buonocore-Buzzelli L, Moss B, et al. Partial efficacy of a VSV-SIV/MVA-SIV vaccine regimen against oral SIV challenge in infant macaques. Vaccine. 2011;29(17):3124–37.

58. Dennis M, Eudailey J, Pollara J, McMillan AS, Cronin KD, Saha PT, et al. Coadministration of CH31 Broadly Neutralizing Antibody Does Not Affect Development of Vaccine-Induced Anti-HIV-1 Envelope Antibody Responses in Infant Rhesus Macaques. Journal of virology. 2019;93(5).

59. Andrews S. FastQC: a quality control tool for high throughput sequence data.. http://www.bioinformatics.babraham.ac.uk/projects/fastqc/.

60. Krueger F. A wrapper tool around Cutadapt and FastQC to consistently apply quality and adapter trimming to FastQ files, with some extra functionality for MspI-digested RRBS-type (Reduced Representation Bisufite-Seq) libraries. http://www.bioinformatics.babraham.ac.uk/projects/trim_galore/.

61. Kersey PJ, Staines DM, Lawson D, Kulesha E, Derwent P, Humphrey JC, et al. Ensembl Genomes: an integrative resource for genome-scale data from non-vertebrate species. Nucleic Acids Res. 2012;40(Database issue):D91–7.

62. Dobin A, Davis CA, Schlesinger F, Drenkow J, Zaleski C, Jha S, et al. STAR: ultrafast universal RNA-seq aligner. Bioinformatics. 2013;29(1):15–21.

63. Liao Y, Smyth GK, and Shi W. featureCounts: an efficient general purpose program for assigning sequence reads to genomic features. Bioinformatics. 2014;30(7):923–30.

64. Love MI, Huber W, and Anders S. Moderated estimation of fold change and dispersion for RNA-seq data with DESeq2. Genome Biol. 2014;15(12):550.

65. Huber W, Carey VJ, Gentleman R, Anders S, Carlson M, Carvalho BS, et al. Orchestrating high-throughput genomic analysis with Bioconductor. Nat Methods. 2015;12(2):115–21.

66. Zhu A, Ibrahim JG, and Love MI. Heavy-tailed prior distributions for sequence count data: removing the noise and preserving large differences. Bioinformatics. 2019;35(12):2084–92.

