## Supplemental for "Distinct upper airway epithelium interferon-stimulated and profibrotic gene expression between adult and infant rhesus macaques infected with SARS-CoV-2"

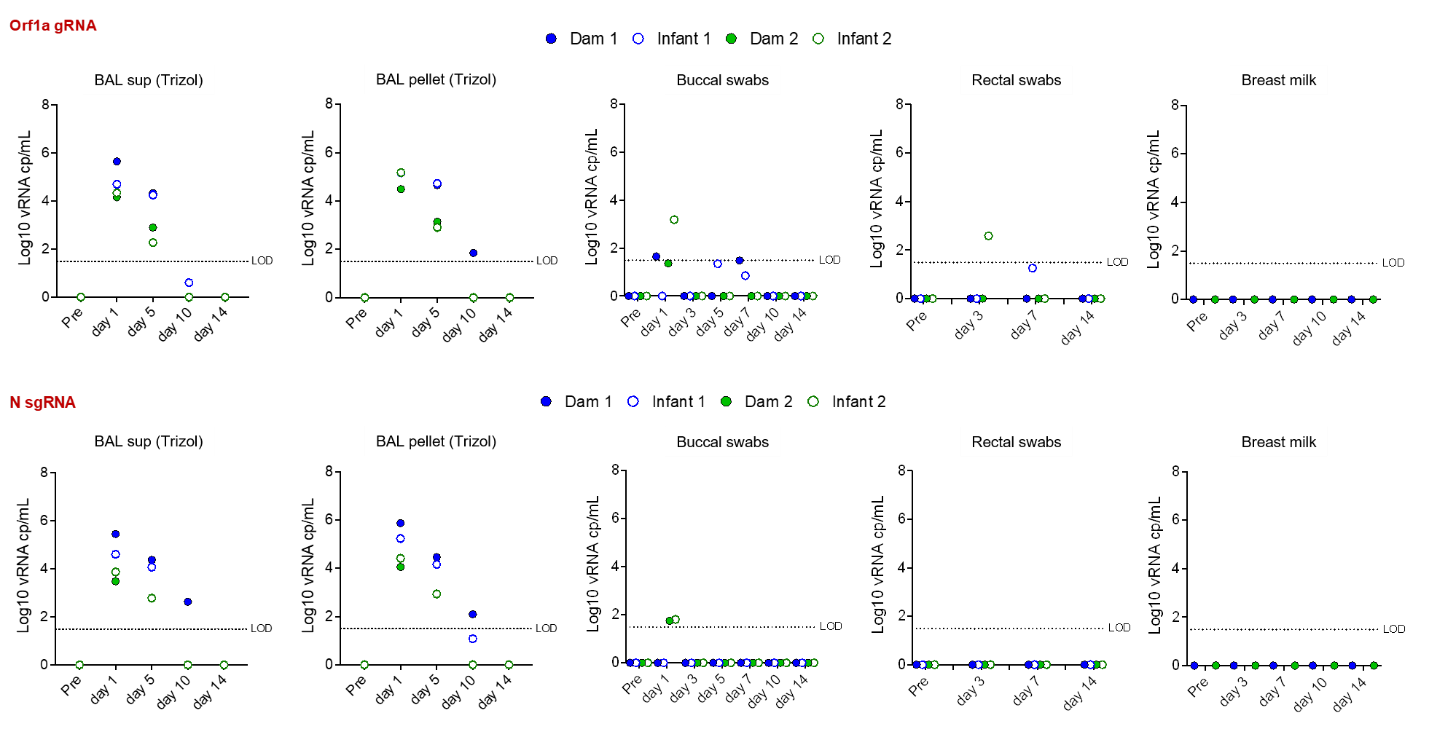

Supplemental Fig. 1

**No significant differences in viral loads between adults and infants in BAL supernatant, BAL pellet, buccal swabs, rectal swabs or breast milk.**

Log_10_ viral RNA copies per mL are reported for bronchoalveolar lavage (BAL) supernatant, BAL pellets, buccal swabs, rectal swabs and breast milk for both the genomic Orf1a gene (top) and subgenomic N gene (bottom) viral RNA.

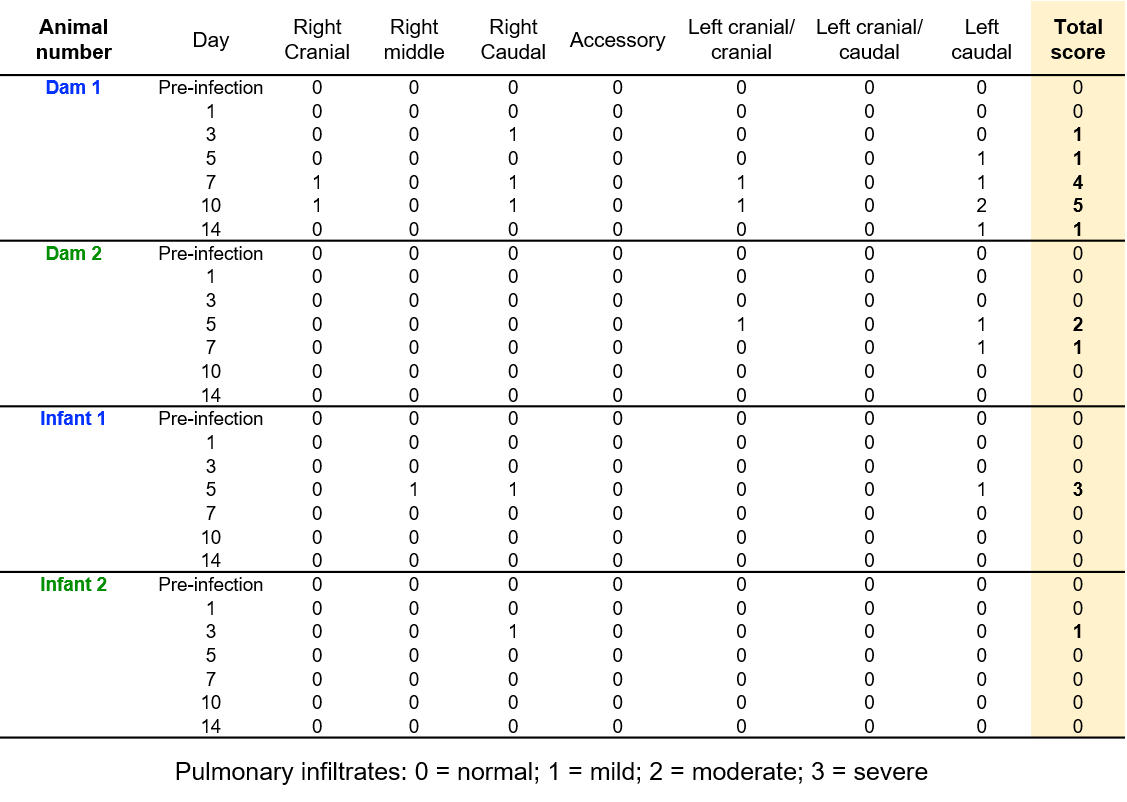

Supplementary Table 1.

Radiographs were scored according to a standard scoring system (0: normal; 1: mild interstitial pulmonary infiltrates; 2: moderate pulmonary infiltrates perhaps with partial cardiac border effacement and small areas of pulmonary consolidation; 3: severe interstitial infiltrates, large areas of pulmonary consolidation, alveolar patterns and air bronchograms). Individual lobes were scored and scores per animal per day were totaled.

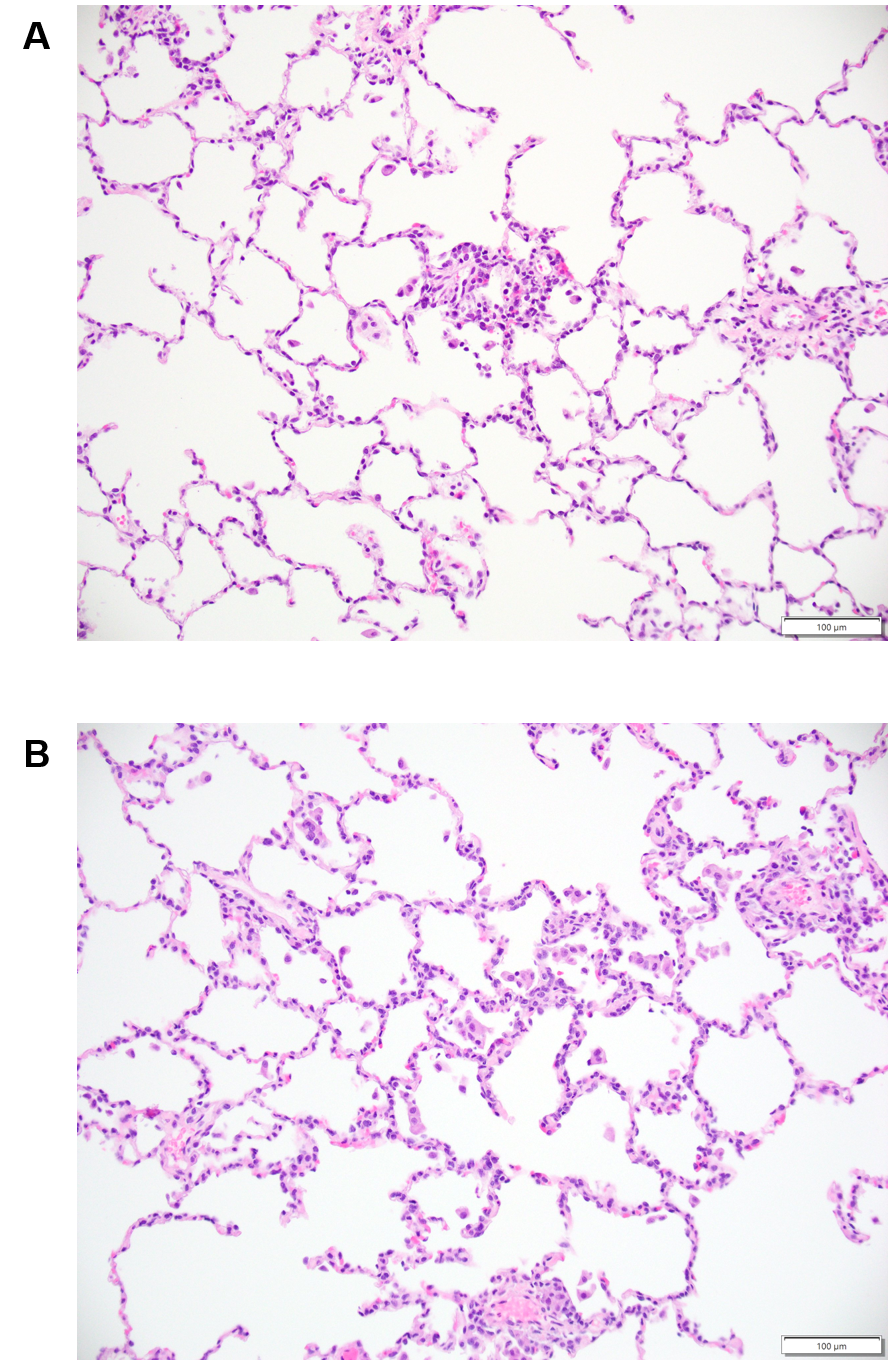

Supplementary Fig. 2.

Interstitial cellularity was evaluated. There was no to minimal interstitial or alveolar cellular infiltrates in either the SARS-CoV infected Dam (B) or infant (A). (H&E, 20x).

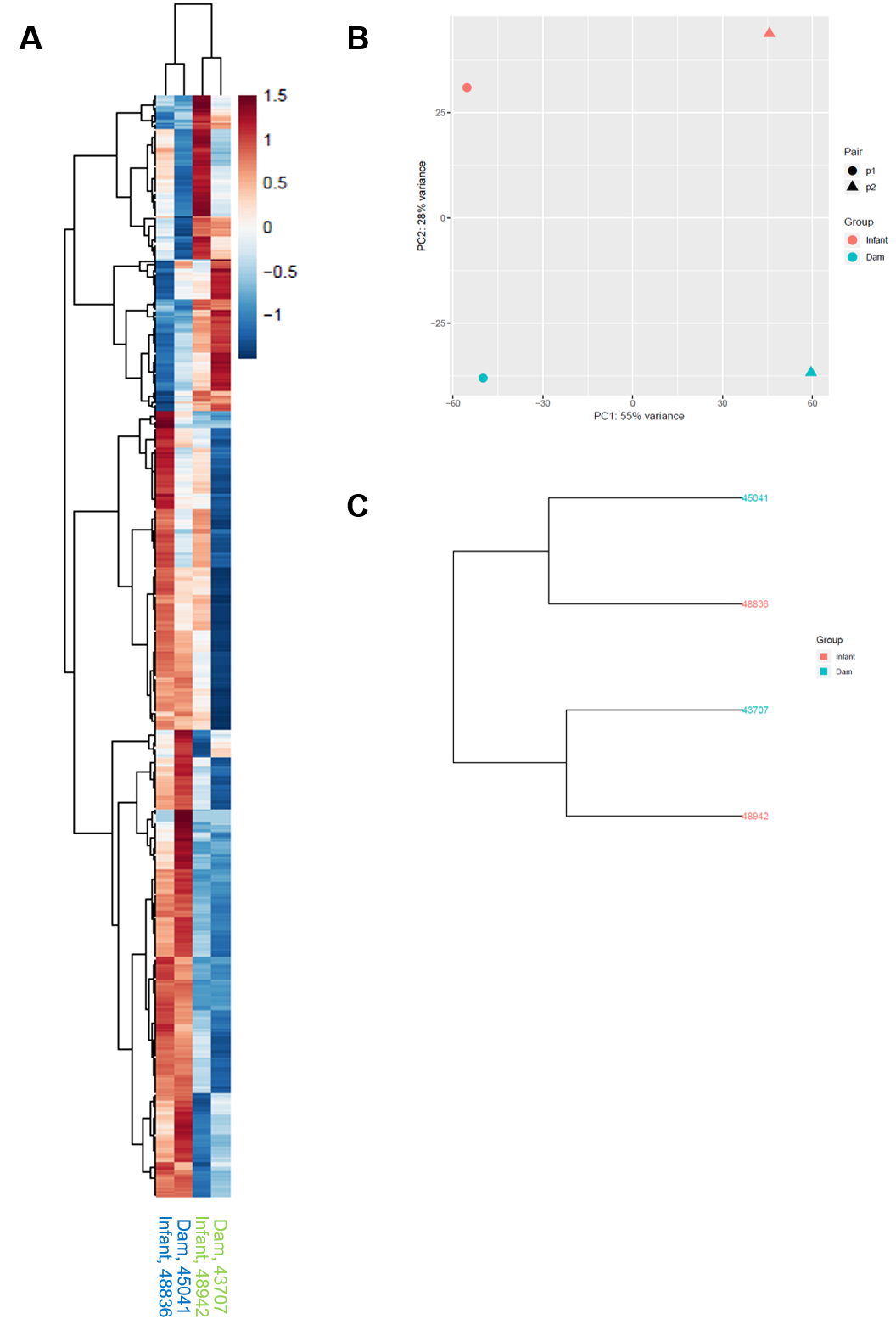

Supplementary Fig. 3.

(A) Heatmap shows expression of differentially expressed genes (FDR ≤ 5%) for each dam and infant comparison. Gene expression has been z-score normalized and the samples and genes are clustered by correlation distance with complete linkage. (B) A principal component analysis (PCA) plot depicts clustering of SARS-CoV-2-infected dam and infant rhesus macaques. (C) Hierarchical clustering based on all genes using a correlation distance with complete linkage. The sample names are colored by condition (infant vs dam).

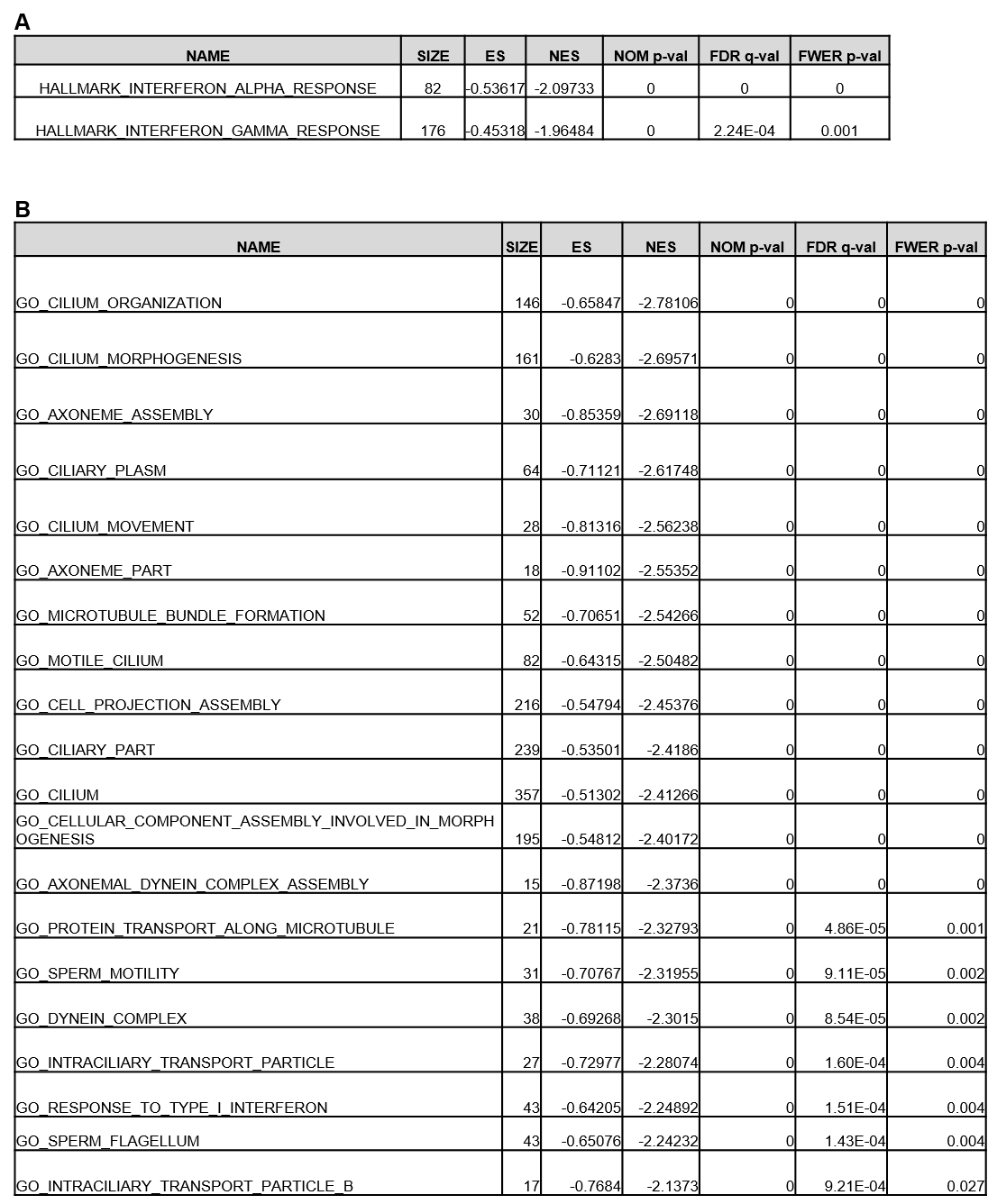

Supplementary Table 2.
(**A**) Hallmark pathways (FWER p-value of ≤0.05) enriched among downregulated genes in SARS-CoV-2 infected adult compared to infant rhesus macaques on day 14 post-infection. (**B**) Gene Ontology (GO) pathways (FWER p-value of ≤0.05) enriched among downregulated genes in SARS-CoV-2 infected adult compared to infant rhesus macaques on day 14 post-infection.

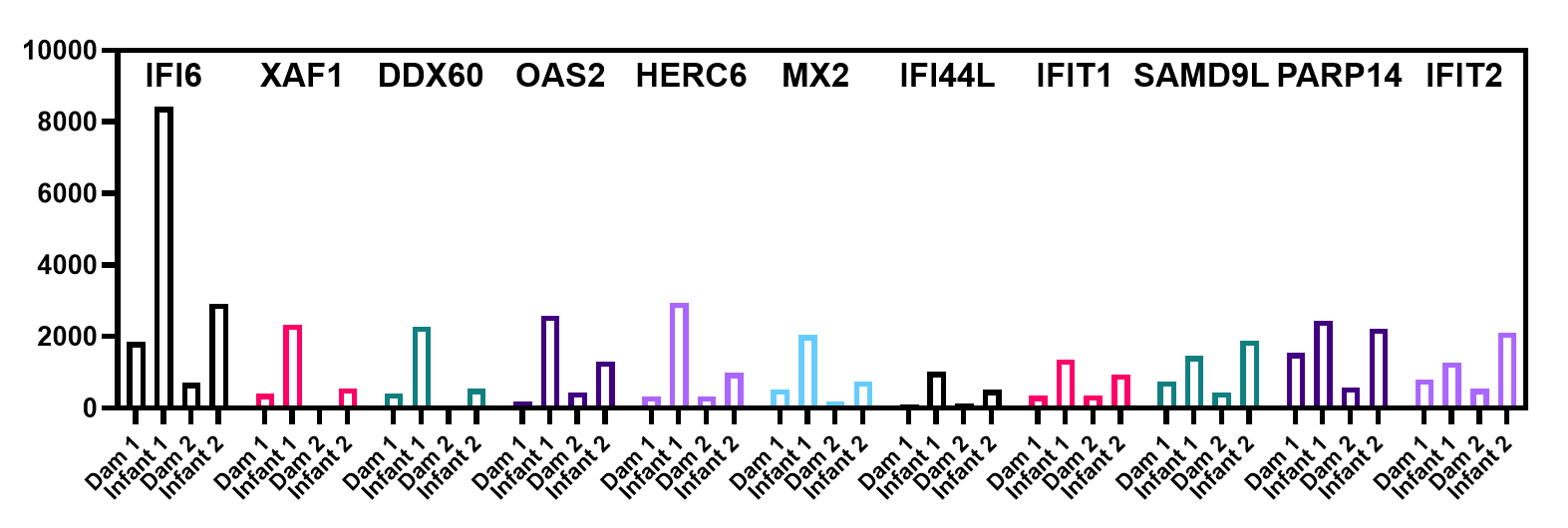

Supplementary Fig. 4.

Normalized expression data from RNA-seq analysis of interferon stimulated genes that are downregulated in adult compared with infant macaques 14 days after SARS-CoV-2 infection.

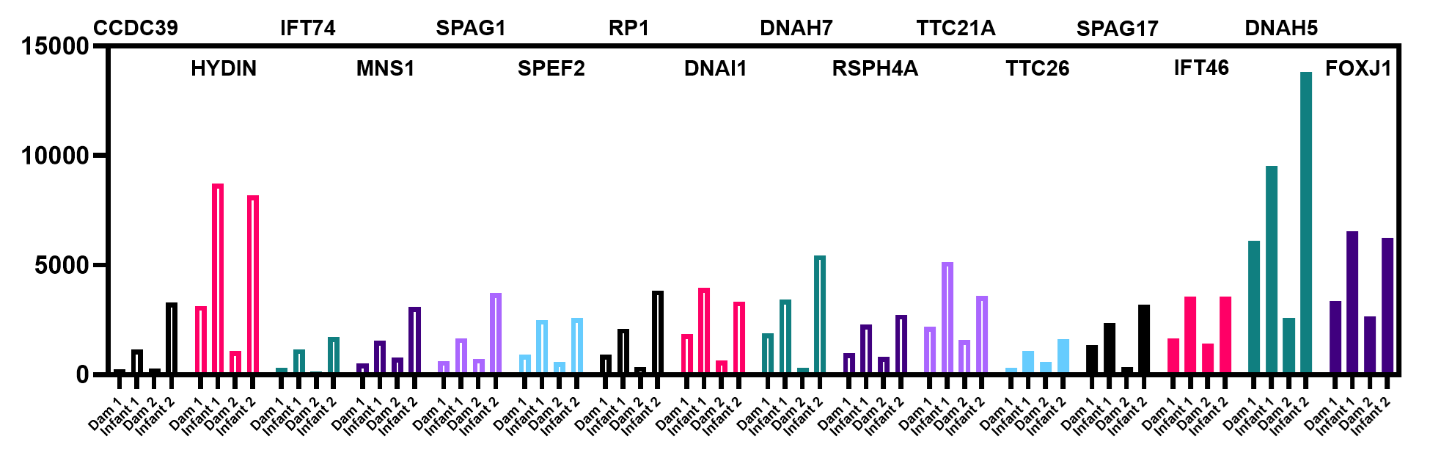

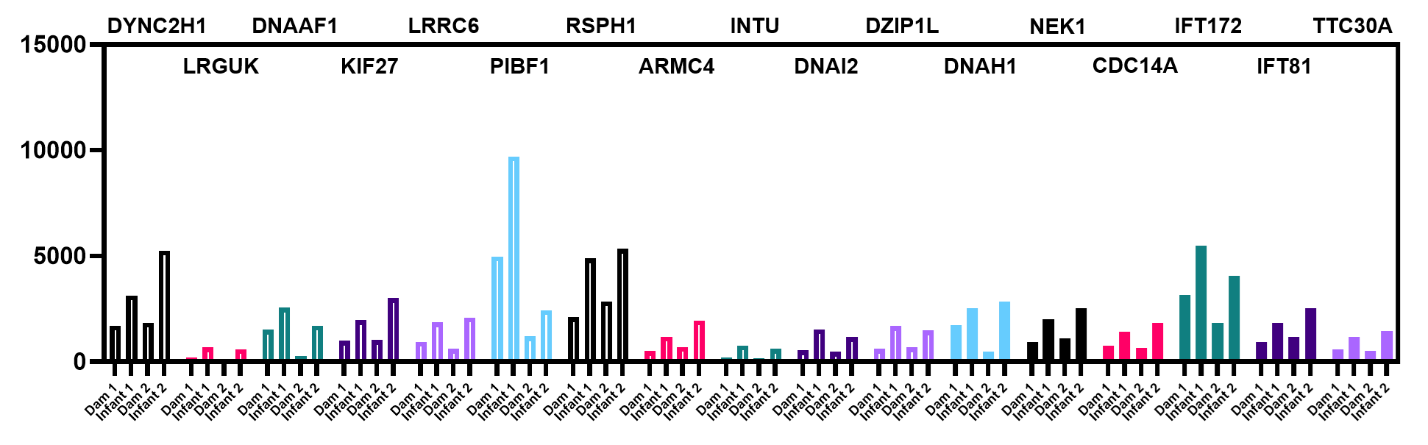

Supplementary Fig. 5.

Normalized expression data from RNA-seq analysis of genes related to cilia structure and function that are downregulated in adult compared with infant macaques 14 days after SARS-CoV-2 infection.

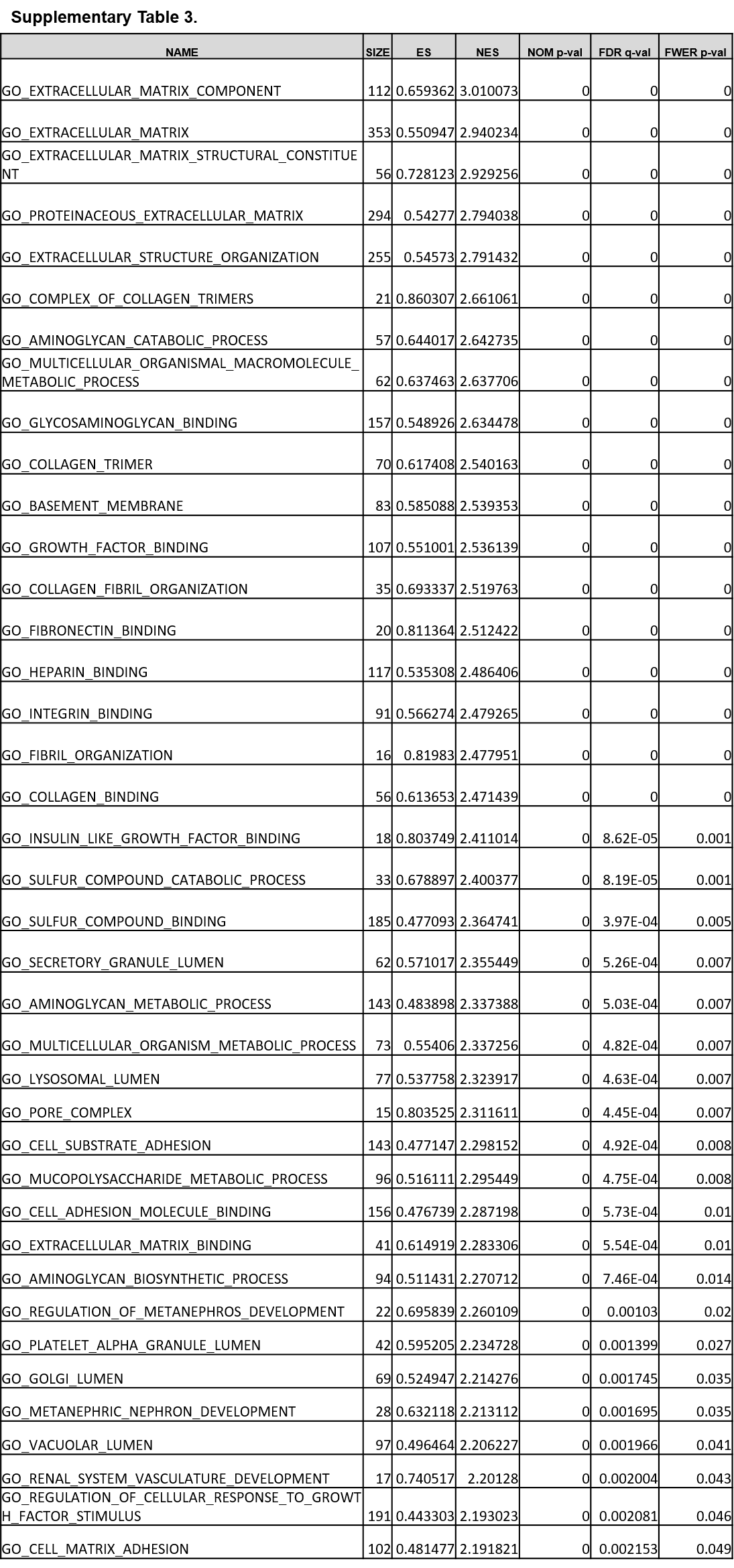

Supplementary Table 3.

Gene Ontology (GO) pathways (FWER p-value of ≤0.05) enriched among upregulated genes in SARS-CoV-2 infected adult compared to infant rhesus macaques on day 14 post-infection.

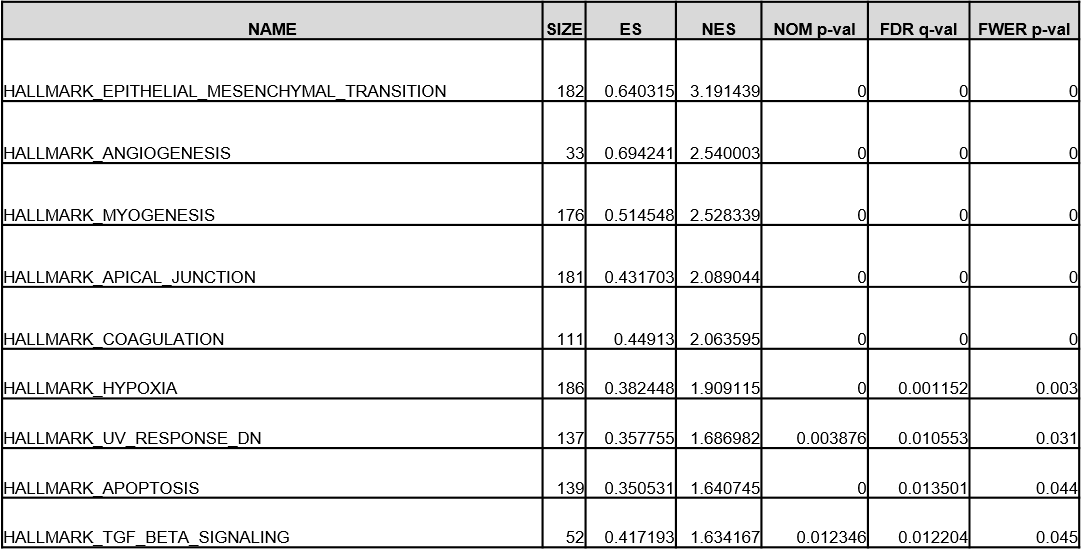

Supplementary Table 4.

Hallmark pathways (FWER p-value of ≤0.05) enriched among upregulated genes in SARS-CoV-2 infected adult compared to infant rhesus macaques on day 14 post-infection.

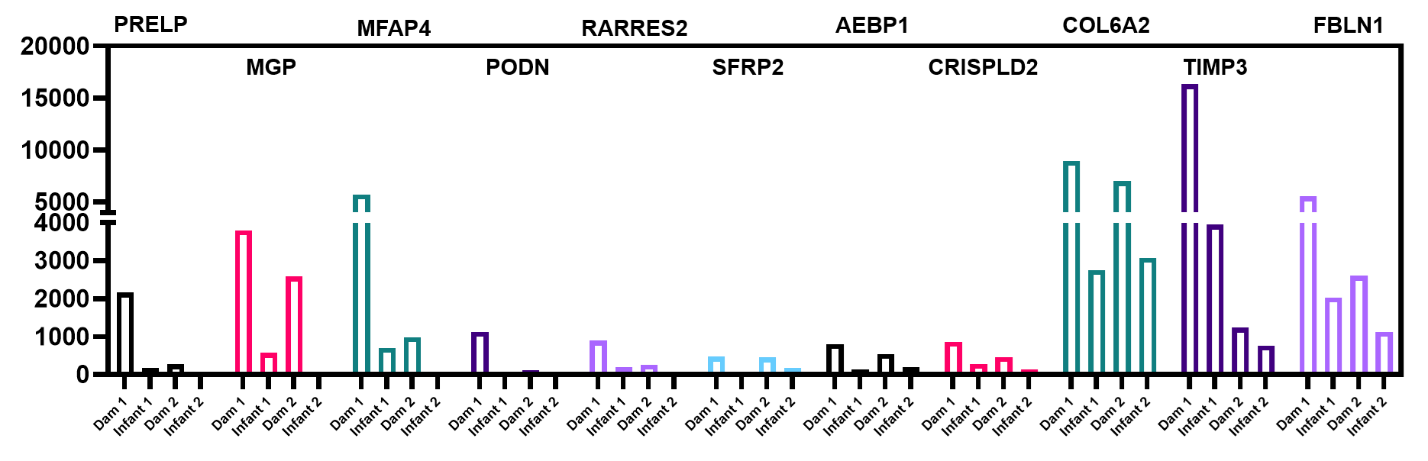

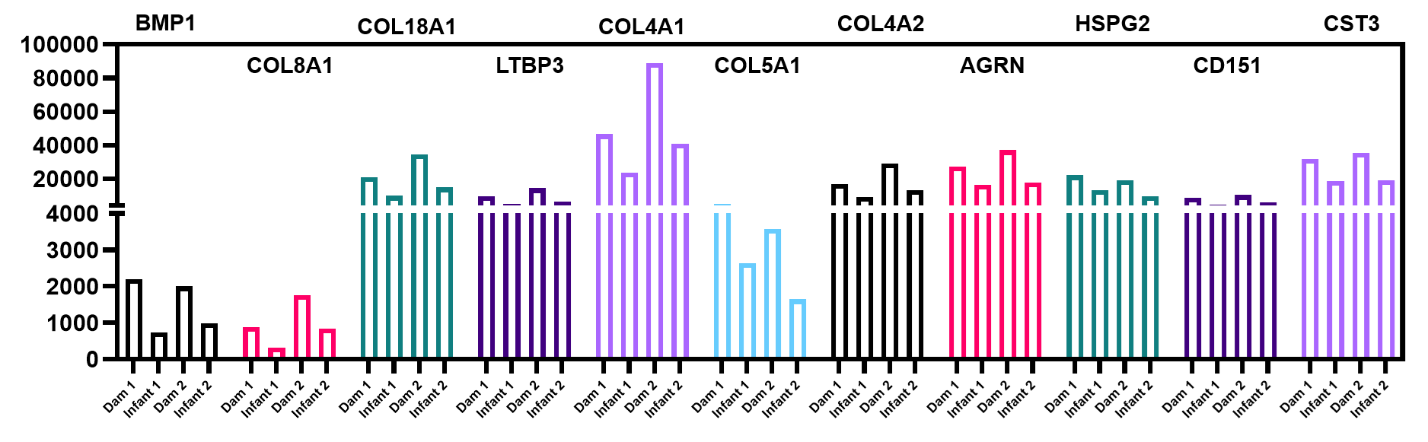

Supplementary Fig. 7.

Normalized expression data from RNA-seq analysis of genes related to extracellular matrix structure and metabolism that are upregulated in adult compared with infant macaques 14 days after SARS-CoV-2 infection.
